# Maternal infection promotes offspring tissue-specific immune fitness

**DOI:** 10.1101/2021.01.13.426542

**Authors:** Ai Ing Lim, Taryn McFadden, Verena M. Link, Seong-Ji Han, Rose-Marie Karlsson, Apollo Stacy, Taylor K. Farley, Oliver J. Harrison, Han-Yu Shih, Heather A. Cameron, Yasmine Belkaid

**Affiliations:** Metaorganism Immunity Section, Laboratory of Host Immunity and Microbiome, National Institute of Allergy and Infectious Diseases, National Institutes of Health, Bethesda, MD 20892, USA; Section on Neuroplasticity, Mood and Anxiety Disorders Program, National Institute of Mental Health, National Institutes of Health, Bethesda, MD, 20892, USA; Postdoctoral Research Associate Training Program, National Institute of General Medical Science, Bethesda, MD, 20892, US; Kennedy Institute of Rheumatology, Nuffield Department of Orthopaedics, Rheumatology and Musculoskeletal Sciences, University of Oxford, Roosevelt Drive, Oxford OX3 7FY, UK; Center for Fundamental Immunology, Benaroya Research Institute, Seattle, WA 98101, USA; Neuro-Immune Regulome Unit, National Eye Institute, National Institutes of Health, Bethesda, MD, 20892, USA; NIAID Microbiome Program, National Institute of Allergy and Infectious Diseases, National Institutes of Health, Bethesda, MD 20892, USA

## Abstract

The mammalian immune system has evolved in the face of microbial exposure. How maternal infection experienced at distinct developmental stages shapes the offspring immune system remains poorly understood. Here we show that during pregnancy, maternally restricted infection can have permanent and tissue-specific impacts on offspring immunity. Mechanistically, maternal IL-6 produced in response to infection can specifically and directly impose epigenetic changes on fetal intestinal epithelial cells. Such imprinting is associated with long-lasting impacts on intestinal immune homeostasis, characterized by enhanced tonic immunity to the microbiota and heightened Th17 responses within the gut, but not at other barrier sites. Furthermore, the offspring from IL-6-exposed dams developed enhanced protective immunity to gastrointestinal infection. Together, this work demonstrates that maternal infection experienced during pregnancy can be coopted by the fetus to promote long-term tissue-specific fitness.

**Summary sentence:** Infection-induced maternal IL-6 impacts offspring epithelial cells, resulting in heightened immunity to the microbiota and pathogens.

---

The immune system responds to infectious challenges in a manner that both controls invading agents and restores tissue homeostasis. One fundamental property of the immune system is its ability to develop memory of previous encounters, resulting in enhanced responsiveness to subsequent challenges. While immunological memory had initially been characterized as a defining property of the adaptive immune system, a growing body of evidence supports the idea that components of the innate system can also be profoundly impacted by inflammatory or infectious encounters—a phenomenon referred to as trained immunity *(1)*. Recent work also uncovered that tissue stem cells can develop memory of previous insults, resulting in accelerated responses to subsequent challenges *(2, 3)*. Innate imprinting is usually transient, lasting a few weeks to months post-exposure, however, parental experiences have been shown to be transferred across generations in invertebrates, and the phenomenon is also suggested to exist in vertebrates *(4-6)*. Education of innate or stem cells by previous inflammatory insults has been associated with epigenetic reprograming *(2, 3, 7)*, though the mechanisms underlying this fundamental property remain largely elusive and represent an active area of investigation. This novel understanding of immunity, as a phenomenon encompassing the whole organism experiences as opposed to specialized cells, provides a framework to understand the enhanced immune fitness observed in animals raised in physiological environments, compared to animals raised under highly controlled settings *(8, 9)*.

The concept of host immune imprinting is of particular interest in the context of pregnancy, as it represents a fundamental developmental window for the immune system. Pregnancy is associated with unique maternal adaptations that serve to maintain tolerance to the fetus *(10)*. At the same time, the mother can contribute to offspring protection against pathogens via transfer of immunomodulatory factors, including maternal antibodies transferred *in utero* and through breastfeeding *(11)*. How infection experienced by the mother during pregnancy shapes offspring immunity in the long-term remains unknown. Indeed, most of our current understanding derives from studies illustrating detrimental outcomes of severe maternal inflammation or infections with pathogens that are able to cross the placental barrier *(12)*. However, mammals and their immune systems evolved in the face of a microbially rich environment, and frequent exposure to pathogens and associated protective immunity represents the norm, rather than the exception. Host survival also requires prioritization of defense strategies that protect the host from tissue damage, in some cases by limiting infection severity without affecting the pathogen load, a phenomenon referred to as disease tolerance *(13, 14)*. As a result, a vast majority of microbial encounters are expected to be associated with mild or no symptoms.

Here we explored the possibility that maternal infection experienced during pregnancy could impact the offspring immune system in the long-term. Based on the fundamental developmental processes occurring *in utero*, we proposed that unique signals experienced at specific developmental windows may imprint the fetus in a discrete and tissue-specific manner.

To assess the impact of maternally restricted infection on offspring immunity, we utilized an attenuated strain of the food-borne pathogen *Yersinia pseudotuberculosis (yopM) (15)*. Timed-pregnant dams were orally infected with *yopM* at gestation day 10.5 (**Fig. 1A**). This specific time point represents a critical developmental stage for the fetal immune system, characterized by the emergence of hematopoietic stem cells and initiation of hematopoiesis *(16, 17)*. Following *yopM* infection, dams experienced a transient weight loss from days 5 to 8 post-infection, but no other symptoms were observed (**fig. S1A**). Timed-pregnant dams transiently shed *yopM* in their stool, and only 20% of pregnant females experienced transient bacterial dissemination to the spleen and the liver (**fig. S1, B and C**). In a manner comparable to non-pregnant mice *(15), yopM* was significantly controlled by day 5 post-infection and cleared by day 9 (**fig. S1B**). No bacteria were detected in the placenta or the fetal liver at any time point tested, and infection did not impact litter sizes (**fig. S1, C and D**). Thus, *yopM* infection represents a model of mild, maternally restricted transient infection.

**Figure 1.**
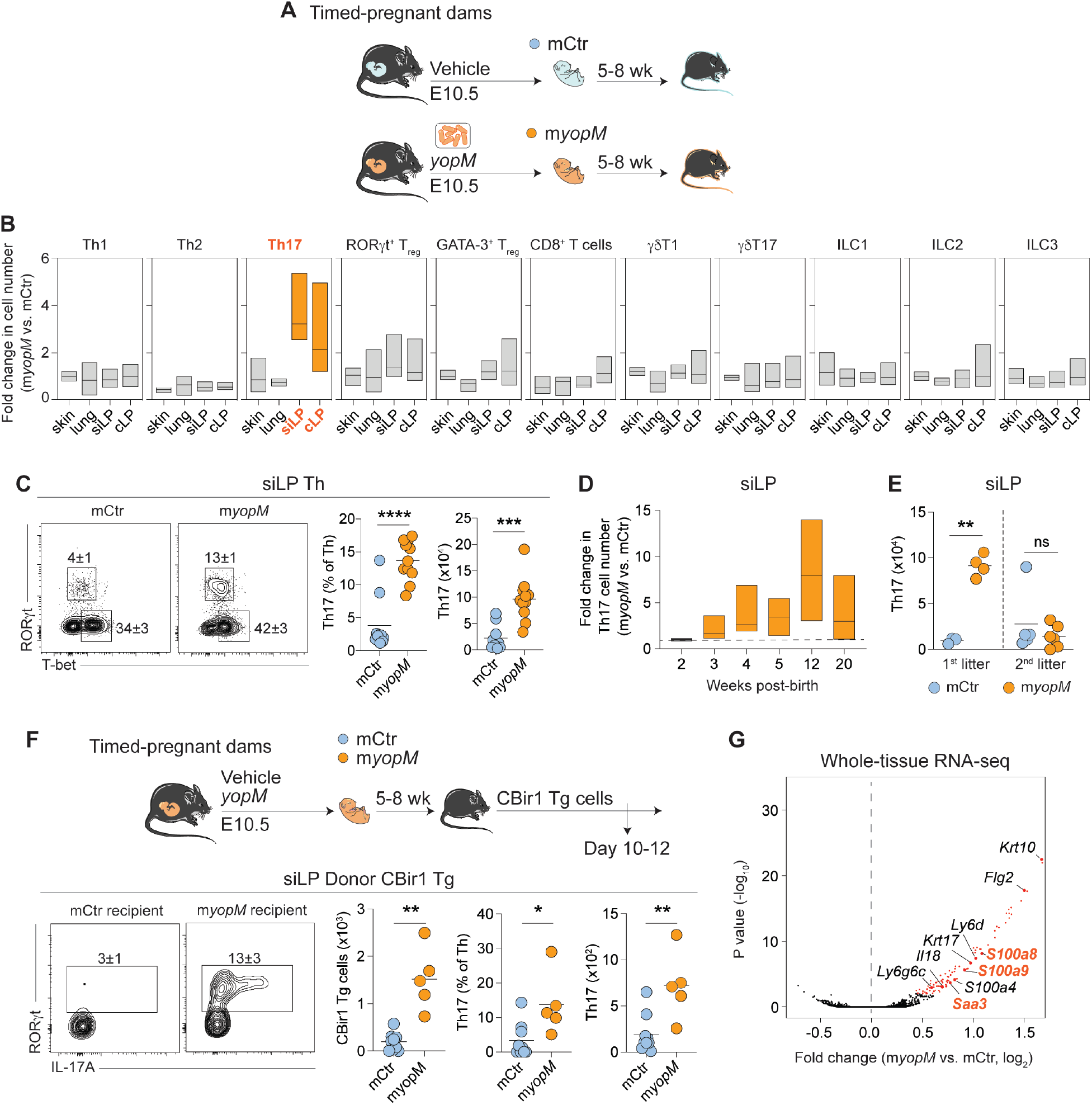
Maternally restricted infection induces tissue-specific alterations to offspring immunity. (**A**) Timed-pregnant dams were orally administered with vehicle (PBS) or *Y. pseudotuberculosis ΔyopM (yopM)* on gestation day 10.5 (E10.5), and their offspring, mCtr or m*yopM*, were analyzed at 5 to 8 weeks old. (**B**) Fold change in cell number of lymphocyte subsets in ear skin, lung, small intestinal lamina propria (siLP) and colonic lamina propria (cLP) of m*yopM* offspring compared to the mean of mCtr offspring. Cells were gated according to the schematic provided in fig. S1E. (**C**) Left: Representative contour plots of transcription factors expressed by offspring siLP T helper (Th) subsets (gated on Live CD45^+^ CD90.2^+^ TCRβ^+^ CD4^+^ Foxp3^-^). Right: Frequency of Th17 (RORγt^+^) among total Th, and total siLP Th17 cell number. (**D**) Fold change in Th17 cell number in the siLP of m*yopM* offspring compared to the mean of mCtr offspring at corresponding ages. (**E**) Total siLP Th17 cell number in the offspring of vehicle-treated dams or dams infected during pregnancy (1^st^) or the offspring from a subsequent pregnancy of the same dams (2^nd^). (**F**) Donor CBir1 Tg cells (CD45.1) were transferred to congenic mCtr or m*yopM* offspring (CD45.2). Donor cells in the siLP were analyzed for transcription factor expression and cytokine production after restimulation at days 10-12 post-transfer. Contour plots were gated on live CD45.1^+^ CD90.2^+^ TCRβ^+^ CD4^+^ Foxp3^-^. (**G**) Whole-tissue RNA-seq of ileum from m*yopM* offspring compared to mCtr offspring. Upregulated genes are denoted in red. (**A-G**) Data are representative of 3 independent experiments, each experiment with 1-2 pregnant dams per group, using 3-5 offspring per pregnancy. Each dot represents an individual mouse. Numbers in representative flow plots indicate mean ± SD. * p < 0.05; ** p < 0.01; *** p < 0.001; **** p < 0.0001; ns, not significant (two-tailed unpaired Student’s t test). See also Figure S1.

We next compared the lymphocyte compartments of the adult offspring delivered by naïve versus previously infected dams. To this end, we characterized the number, frequency and phenotype of T cells (αβ and γδ) and innate lymphoid cells (ILCs) at various barrier tissues, including the skin, the lung and the small intestinal lamina propria (siLP) and colonic lamina propria (cLP) of 5 to 8-week-old offspring (**fig. S1E**). Maternal infection during pregnancy was associated with a significant and selective increase in the number of RORγt-expressing CD4^+^ T cells (Th17) in the small intestinal and colonic lamina propria of the offspring, compared to the animals delivered from naïve dams (**Fig. 1, B and C**). Notably, the increase in Th17 cell number was restricted to the gut compartment and was not detected at other barrier sites assessed (**Fig. 1B**). Furthermore, no differences were observed in the number of other lymphocyte subsets, including T-bet-expressing CD4^+^ T cells (Th1), GATA-3-expressing CD4^+^ T cells (Th2), Foxp3-expressing CD4^+^ regulatory T cells (RORγt^+^ T_reg_ and GATA-3^+^ T_reg_), CD8^+^ T cells, γδ T cells (γδT1 and γδT17) and innate lymphoid cells (ILC1, 2 and 3) in all the compartments assessed (**Fig. 1B and fig. S1F**). Increased accumulation of Th17 cells within the siLP was detectable post-weaning (at 3 weeks) and persisted throughout adulthood (up to 20 weeks) (**Fig. 1D and fig. S1G**). The impact of infection on the offspring gut immune system was not observed in the offspring from a subsequent litters from the same dams, supporting the idea that the ability of the dams to enhance offspring immunity was not caused by long-term maternal imprinting (**Fig. 1E**). Thus, a transient infection during pregnancy can impose long-term, tissue-specific alterations in the offspring’s immunity.

We next assessed whether this phenomenon was T cell-intrinsic or caused by an altered tissue environment. To this end, we evaluated the ability of the offspring delivered by control or previously infected dams to mount *de novo* responses to the microbiota, utilizing T cells specific for a defined commensal-derived antigen (CBir1 Tg cells) *(18)*. Transgenic CBir1 T cells were transferred into the offspring of naïve or previously infected dams (**Fig. 1F**). Consistent with previous studies *(18, 19)*, a small number of CBir1 Tg cells accumulated in the siLP of control mice where they remained largely undifferentiated (**Fig. 1F**). In contrast, a higher number of CBir1 Tg cells were found in the siLP of the offspring delivered by previously infected dams, with a significant fraction of those differentiated towards a Th17 phenotype (**Fig. 1F**). Thus, maternal infection impacted the offspring intestinal milieu in a way that enhanced reactivity toward the microbiota and promoted Th17 responses. In support of this observation, whole-tissue RNA sequencing (RNA-seq) revealed alterations in the intestinal microenvironment, including upregulation of genes encoding antimicrobial peptides *(S100a8* and *S100a9)* and serum amyloid A *(Saa3)* (**Fig. 1G**), factors previously shown to promote local licensing of Th17 cells *(20)*.

Infection can impact the composition, localization and function of the microbiota *(21, 22)*. As such, we next explored the possibility that these enhanced Th17 responses could be associated with maternally acquired microbiota. To this end, we compared the gut microbiota of the offspring delivered by previously infected versus control dams, utilizing 16S rRNA gene sequencing (**Fig. 2A**). The microbiota of the offspring delivered by infected and naïve dams clustered closely to each other, as demonstrated by Bray-Curtis principal coordinates analysis (**Fig. 2B**). Though no significant shifts occurred at the phylum level, a small number of species belonging to the *Bacteroidales* and *Coriobacteriales* families were increased, while a few species belonging to the *Clostridiales* and *Erysipelotrichales* families were decreased, in the offspring of previously infected dams (**Fig. 2, C and D**). To assess the potential contribution of the altered microbiota to offspring immunity, newborns delivered by control and previously infected dams were cross-fostered until weaning (**Fig. 2E**). Offspring delivered by previously infected dams and fostered by control dams harbored a number of Th17 cells within the siLP at a level comparable to offspring delivered and fostered by previously infected dams (**Fig. 2, F, G, H and I**). On the other hand, offspring delivered by control and fostered by previously infected dams did not exhibit enhanced Th17 cell number within the siLP (**Fig. 2, F, G, H and I**). These data support the idea that the heightened intestinal Th17 responses caused by maternal infection were acquired *in utero*, and did not result from the transfer of altered maternal microbiota following birth or during lactation.

**Figure 2.**
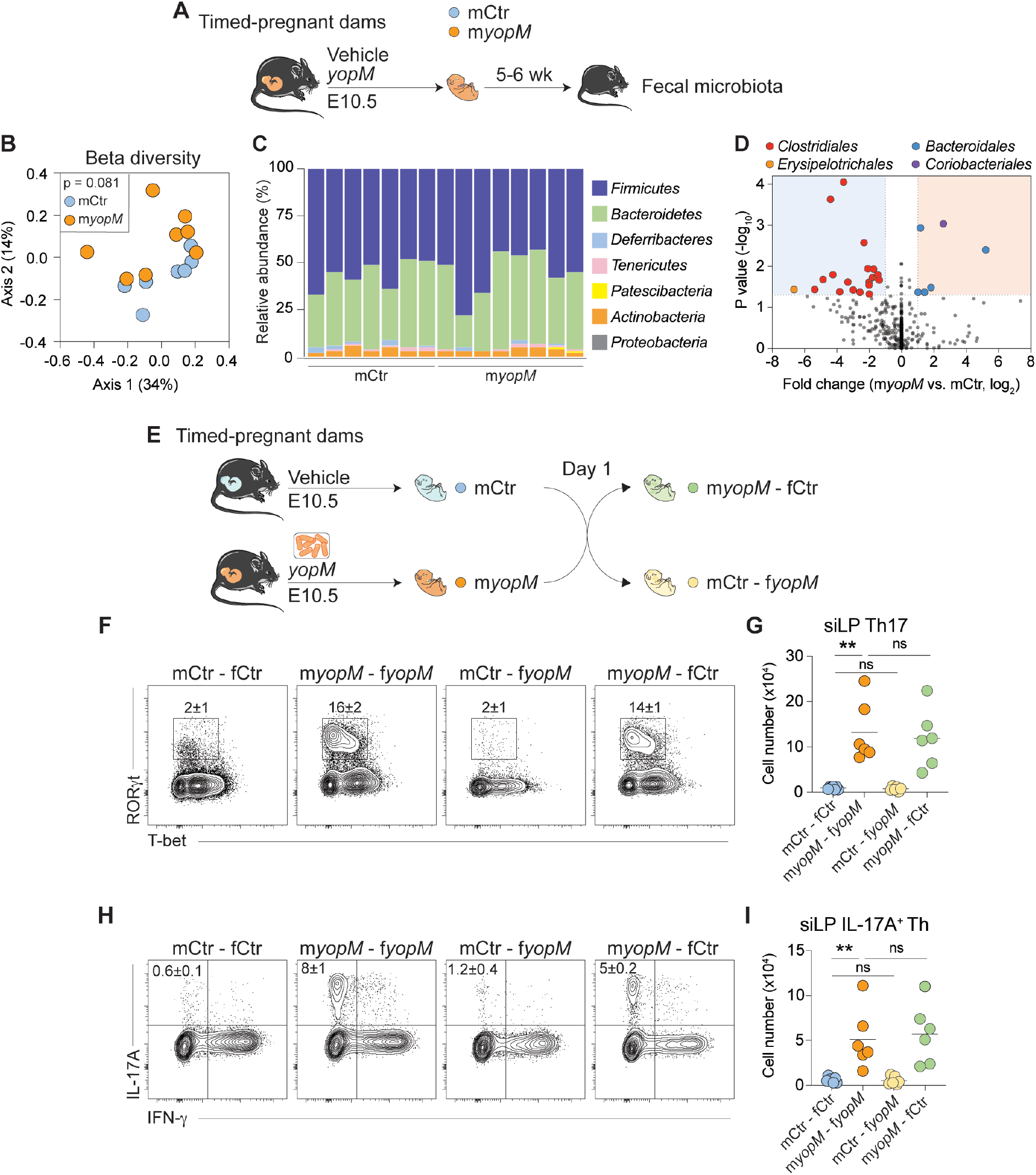
Maternally restricted infection alters offspring intestinal immunity *in utero*, independently of the microbiota. (**A**) Timed-pregnant dams were orally administered with vehicle or *yopM*, and fecal microbiota from their offspring, mCtr or m*yopM*, were profiled at 5 to 6 weeks old by 16S rRNA gene sequencing. (**B**) Principal coordinates analysis (PCoA) plot displaying the Bray-Curtis distances between the fecal 16S profiles of mCtr and m*yopM* offspring. Percentages represent the variance corresponding to each PC. Significance was determined by PERMANOVA. (**C**) Relative abundance of phyla in the mCtr and m*yopM* fecal microbiota. (**D**) Volcano plot of fecal microbiota families. (**E**) Cross-fostering experiment: Within 24 hours after birth, offspring delivered by vehicle-treated dams were transferred to previously-infected dams (mCtr-f*yopM)* or *vice versa* (m*yopM*-fCtr). Offspring were cross-fostered until weaning and analyzed at 5 to 8 weeks old. In the control groups, the offspring were raised by their own dams (mCtr-fCtr, m*yopM*-f*yopM)*. (**F**) Representative contour plots of transcription factors expression by Th subsets in the offspring siLP. (**G**) Total siLP Th17 (RORγt^+^) cell number. (**H**) Representative contour plots of cytokine production by Th after restimulation. (**I**) Total cell number of IL-17A-producing Th cells. (**A-I**) All flow plots were gated on live CD45^+^ CD90.2^+^ TCRβ^+^ CD4^+^ Foxp3^-^. Numbers in representative flow plots indicate mean ± SD. Data are representative of 3 independent experiments, each experiment with 1 pregnant dam per group, using 3-6 offspring per pregnancy. Each dot represents an individual mouse. ** p < 0.01; *** p < 0.001; ns, not significant (one-way ANOVA multiple-comparison test).

Soluble factors including cytokines can cross the placental barrier (23, 24). To assess a potential role for maternal soluble factors, we transferred sera from infected dams to naïve dams at gestation day 13.5 and 15.5 (Fig. 3A). We confirmed that no bacteria were transferred, and that the recipient dams did not mount an immune response against yopM (fig. S2, A and B). Injection of sera from previously infected dams was sufficient to increase the offspring intestinal Th17 cell number, in a manner comparable to that observed after maternal yopM infection (Fig. 3A). Assessment of cytokine and chemokine levels in the sera of yopM-infected pregnant dams at different time points post-infection revealed that 4 cytokines were significantly enriched, including IL-6, IFN-g, G-CSF and TNF-a (**Fig. 3B**). We next tested the potential role of individual cytokines in remodeling the offspring immunity by injecting each of these 4 cytokine candidates to naïve timed-pregnant dams. A single injection of IL-6—but not IFN-g, G-CSF or TNF-a—to naïve pregnant dams was sufficient to increase the Th17 cell number in the siLP of the offspring (fig. S2C). The induction of intestinal Th17 cells in the progeny caused by IL-6 injection was dose-dependent (**Fig. 3C**). Of interest, a previous study showed that IL-6 enhanced IL-17A production in dams the production of which promoted the development of an autism-like disorder in offspring *(25)*. However, in our present study, the enhanced IL-6 level during pregnancy did not cause any detectable impairment in social preference (**fig. S2D**). One notable difference in experimental settings was that our study was conducted in mice devoid of segmented filamentous bacteria (SFB), a group of bacteria previously implicated in altered behaviors in progeny *(26)*.

**Figure 3.**
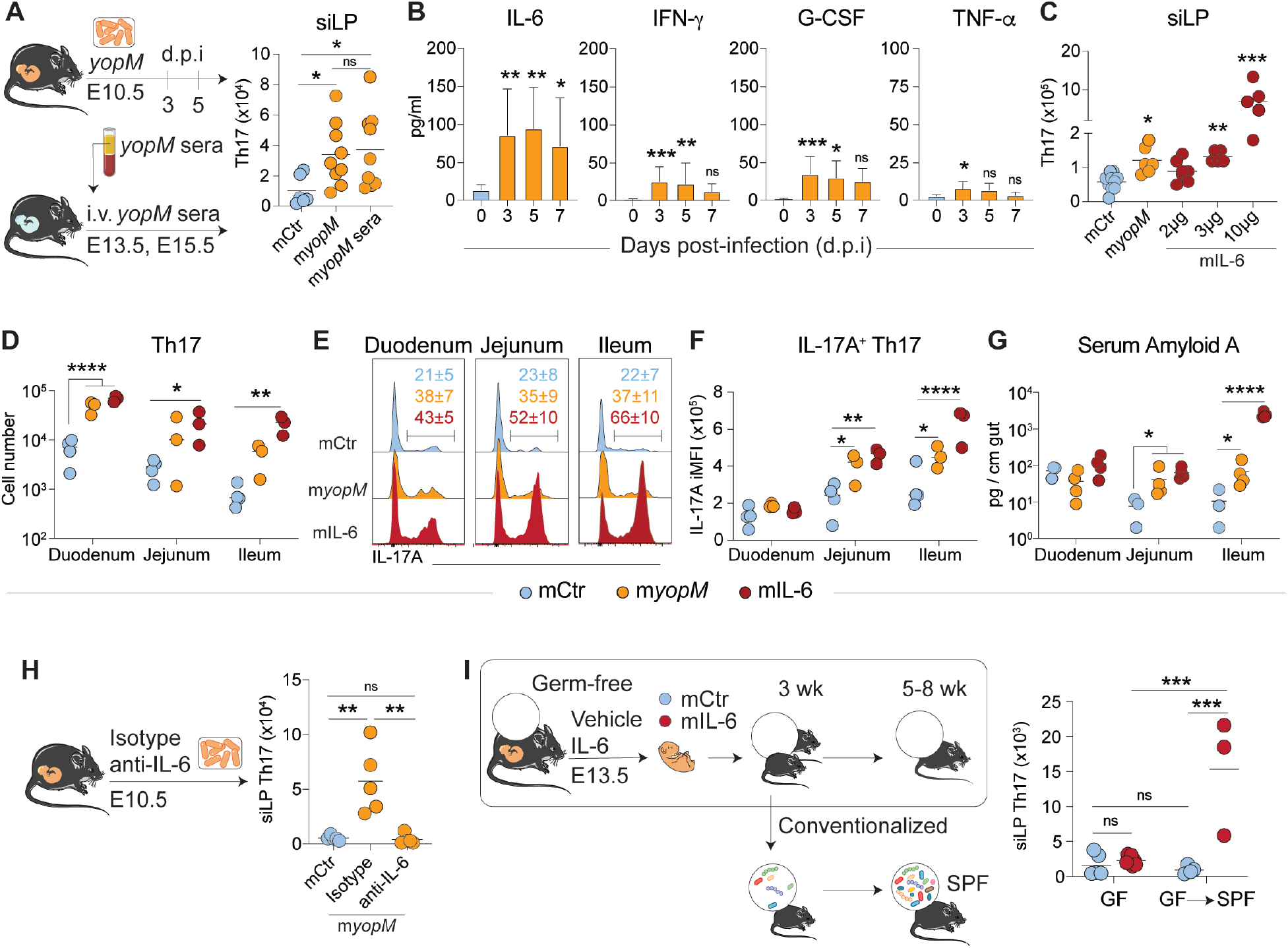
Increasing maternal IL-6 levels during pregnancy is sufficient to enhance offspring intestinal Th17 and responsiveness to the microbiota. (**A**) Left: Timed-pregnant dams were infected with *yopM*, and sera collected at 3 and 5 days post-infection (d.p.i) were transferred intravenously (i.v.) to naïve pregnant dams at E13.5 and E15.5. Right: Total Th17 cell number in the siLP of 6-week-old offspring. (**B**) Timed-pregnant dams were orally infected with *yopM* at E10.5, sera collected before infection or at 3, 5, or 7 d.p.i were assayed for cytokine and chemokine levels. (**C**) Timed-pregnant dams were orally administered with vehicle or *yopM* or i.v. injected with indicated doses of IL-6, and their offspring, mCtr, m*yopM* or mIL-6, were analyzed at 6 weeks old for total siLP Th17 cell number. (**D-G**) siLP sub-compartments from 6-week-old offspring delivered by mCtr, m*yopM* or dams i.v. injected with 10 μg IL-6 (mIL-6) were analyzed for (**D**) total Th17 cell number, (**E**) frequency of IL-17A-producing Th17, (**F**) integrated mean fluorescence intensity (iMFI) of IL-17A production by Th17 (iMFI = (IL-17A MFI) x (frequency of IL-17A-producing Th17)) and (**G**) serum amyloid A level. (**H**) Left: *yopM*-infected dams were pre-treated with 1 mg of anti-IL-6 monoclonal antibody or the isotype control 2 hours prior to infection. Right: Total Th17 cell number in the siLP of 6-week-old offspring. (**I**) Left: Germ-free (GF) timed-pregnant dams were i.v. injected with vehicle or 10 μg of IL-6 at E13.5, and their offspring, mCtr or mIL-6, were maintained under GF conditions or conventionalized in a specific-pathogen-free (SPF) facility at 3 weeks old. Right: Total Th17 cell number in the siLP of 6-week-old offspring. Data are representative of 3 independent experiments, each experiment with 1-2 pregnant dams per group, using 3-5 offspring per pregnancy. Each dot represents an individual mouse. * p < 0.05; ** p < 0.01; *** p < 0.001; **** p<0.0001; ns, not significant (ANOVA multiple-comparison test, one-way in A, B, C, H and two-way in D, F, G, I). See also Figure S2.

As observed in the context of maternal infection, IL-6 injection significantly increased the number of Th17 cells in all compartments of the offspring small intestine (**Fig. 3D**). Furthermore, IL-6 injection was associated with a significant increase in the level of IL-17A production by Th17 in the jejunum and the ileum (**Fig. 3, E and F**). In agreement with enhanced IL-17A production *(20)*, both maternal infection and IL-6 injection enhanced serum amyloid A production in the jejunum and the ileum (**Fig. 3G**). IL-6 injection also induced a small increase in the number of CD8^+^ T cells in the duodenum (**fig. S2E**). Thus, IL-6 injection to dams was sufficient to alter offspring intestinal immunity. To address the possibility that IL-6 may also be necessary for the impact of maternal *yopM* infection on offspring immunity, pregnant dams were treated with a neutralizing IL-6 antibody prior to *yopM* infection (**Fig. 3H**). Such treatment did not affect the ability of the pregnant dams to control the infection (**fig. S2F**). Notably, IL-6 neutralization during maternal infection significantly reduced Th17 cell accumulation in the siLP of the offspring (**Fig. 3H**). Collectively, our findings support the idea that increased level of maternal IL-6, elicited by transient infection during pregnancy, altered offspring intestinal immunity.

While a shift in the maternally acquired microbiota was not responsible for altered offspring immunity (**Fig. 2, D and G**), we assessed whether the maternal microbiota was required during pregnancy and/or post-delivery for the remodeling of the offspring immune response. To address this point, germ-free pregnant dams were injected with IL-6 or vehicle control. IL-6 injection did not cause enhanced Th17 cell accumulation in the offspring siLP if the offspring were maintained under germ-free condition (**Fig. 3I**). On the other hand, reintroduction of the microbiota (conventionalization) post-weaning was associated with enhanced Th17 cell number in the offspring from IL-6-injected, but not in those from control dams (**Fig. 3I**). Thus, prenatal establishment and postnatal maintenance of IL-6-mediated tissue imprinting is independent of the maternal microbiota but allows the offspring to mount enhanced Th17 responses to postnatal exposure to the microbiota.

Responses to IL-6 signaling are mediated by the IL-6 receptor IL-6R*α* (also known as CD126), the glycoprotein 130 complex and the JAK/STAT3 pathway *(27)*. IL-6 has also been shown to cross both human and murine placenta *(28, 29)*. To investigate the impact of IL-6 on the fetal intestine, we first assessed expression of IL-6R*α* in fetal intestinal epithelial and hematopoietic cells. We found that IL-6R*α* was expressed at an intermediate level in all fetal intestinal epithelial cells (IEC, EpCAM^+^) and at a high level in a fraction of hematopoietic cells (CD45^+^) (**Fig. 4, A and B**). To identify cellular compartments that can respond to IL-6, we evaluated STAT3 phosphorylation at day 3 post-*yopM* infection or post-IL-6 injection. A small fraction of IECs expressed phosphorylated STAT3 (pSTAT3) at steady state, supporting the idea that the JAK/STAT3 pathway may already be constitutively engaged during epithelial cell development (**Fig. 4C**). At day 3 post maternal infection or IL-6 injection, pSTAT3 level was further increased in fetal IECs, but not in hematopoietic cells (**Fig. 4C**).

**Figure 4.**
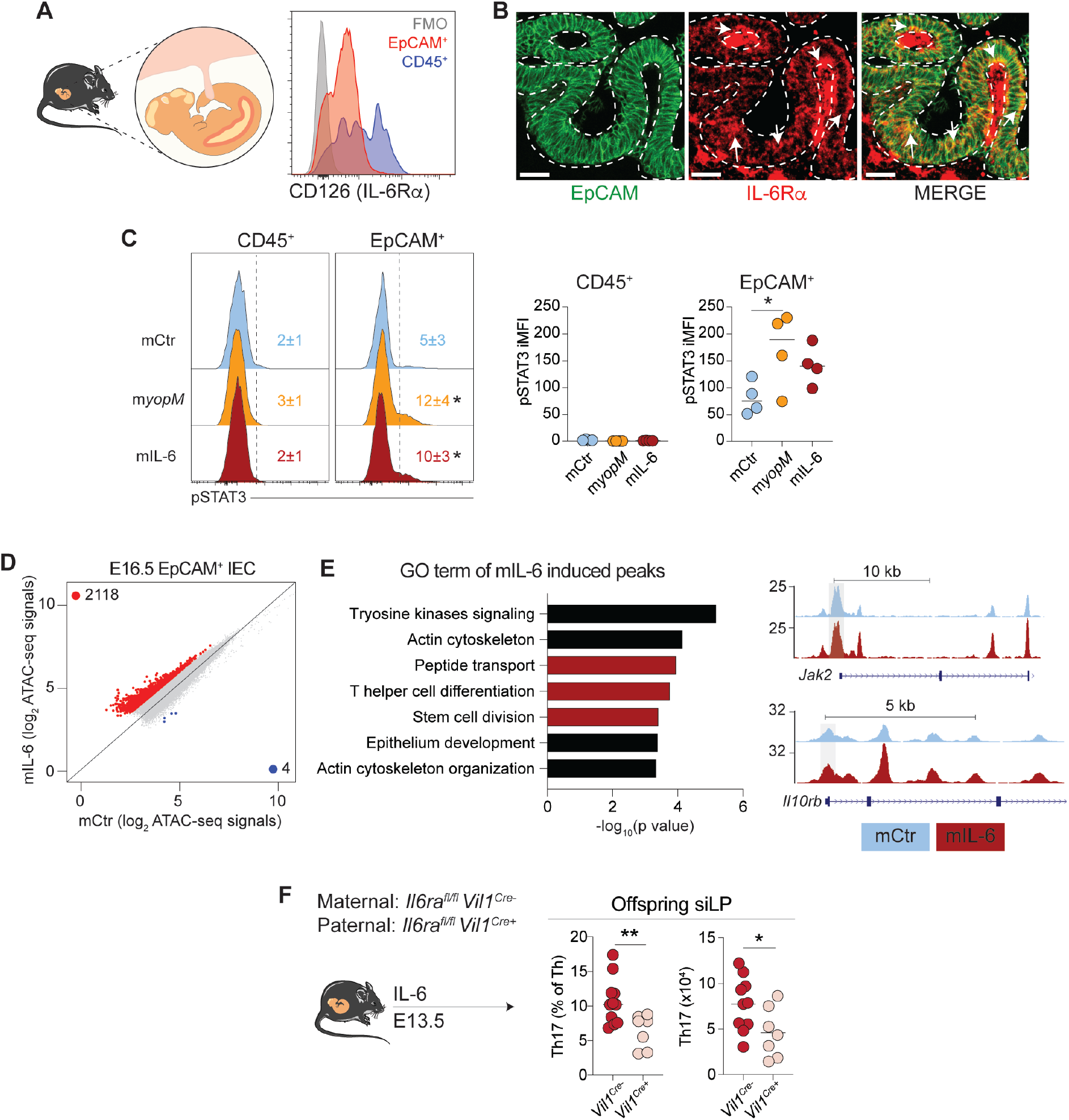
Increasing maternal IL-6 levels during pregnancy alters the epigenome of fetal intestinal epithelial cells. (**A**) Left: Fetal intestines were assessed for IL-6R*α* (CD126) expression. Right: Representative histogram showing IL-6R*α* expression by fetal intestinal epithelial cells (IEC, CD45^-^ EpCAM^+^) and hematopoietic cells (CD45^+^) at E15.5. FMO: fluorescence minus one control. (**B**) Representative micrograph of fetal intestines stained for EpCAM and IL-6R*α* at E15.5. Arrows point to IL-6R*α* expressed by EpCAM^+^ IEC (denoted by dashed line). Scale bars: 30 μm. (**C**) Left: Representative histogram of pSTAT3 expression by fetal hematopoietic cells or IEC from dams at 3 days post-treatment with vehicle (mCtr), *yopM* (m*yopM)*, or 10 μg IL-6 (mIL-6). Right: Integrated MFI (iMFI) of pSTAT3 in hematopoietic cells (CD45^+^) or IECs (EpCAM^+^). (**D-E**) ATAC-seq data of fetal IEC from dams 3 days post-treated with vehicle (mCtr) or i.v. injected with 10 μg of IL-6-injected (mIL-6). (**D**) Scatter plot of all accessible chromatin regions. Red and blue dots correspond to regions that were differentially accessible by >2 fold in the IEC of mIL-6 fetuses and mCtr fetuses, respectively. (**E**) Left: Top GO terms that were enriched in the differentially accessible promoter regions (peaks within 3kb of the next transcription start site, fold change > 2) of mIL-6 fetuses compared to mCtr fetuses. Right: UCSC Genome Browser snapshots of genes of interest. Promoters open in mIL-6 offspring are highlighted in gray boxes. (**F**) *Il6ra* was deleted from IEC by crossing *Il6ra*^fl/fl^ *Vil1*^*Cre-*^ females to *Il6ra*^fl/fl^ *Vil1*^*Cre+*^ males. Dams were injected with 10 μg of IL-6 and offspring were analyzed at 5 to 8 weeks old for siLP Th17 cells. (**A-F**) Data are representative of 3 independent experiments (A-C, F), each experiment with 1-2 pregnant dams per group, or 1 experiment (D-E) with 3 pregnant dams per group, using 4-7 offspring per pregnancy. * p < 0.05; ** p < 0.01 (two-tailed unpaired Student’s t test in F or one-way ANOVA multiple-comparison test in C). See also Figure S3 and S4.

To test the possibility that IECs might be the direct target tissue imprinted by maternal IL-6 during infection, we characterized the epigenetic and transcriptional landscape of fetal IECs from control and IL-6-injected dams, utilizing an assay for transposase-accessible chromatin with high-throughput sequencing (ATAC-seq) and RNA-seq (**fig. S3A**). Genes important for IEC identity (including *Epcam, Vil1, Cdx1, Cdx2) (30, 31)* were equally accessible in fetal IEC from control and IL-6-injected dams (**fig. S3B**). Open chromatin regions in fetal IECs were enriched in transcription factor binding motifs that were previously implicated in intestinal development (CDX1/2), IEC differentiation (GATA, HNF4G) and epithelial cell identity establishment (KLF) (**fig. S3C**) *(32)*. Injection of IL-6 during pregnancy substantially increased fetal IEC chromatin accessibility, with 2118 regions uniquely open (fold change greater than 2), compared to only 4 regions in the IECs from fetuses from control dams (**Fig. 4D**). The differentially accessible promoter regions revealed enrichment of genes associated with intestinal physiological functions, including peptide transport, stem cell division, epithelium development and T helper cell differentiation (e.g. *Jak2, Il10rb)* (**Fig. 4E**). On the other hand, transcriptomic analysis showed minor alterations in fetal IECs (**fig. S3D**). Thus, exposure to IL-6 during development increases global chromatin accessibility of fetal IECs, with minor impacts on their transcriptional landscape.

To further assess the contribution of IL-6 signaling on the ability of fetal IECs to regulate intestinal immunity, we specifically deleted the IL-6 receptor gene *Il6ra* from IECs (**Fig. 4F**). To this end, we utilized a breeding scheme allowing IL-6 signaling to be intact in the dams *(Il6ra*^*fl/fl*^ *Vil1*^*Cre-*^) and for IECs of the offspring to be either *Il6ra* sufficient *(Il6ra*^*fl/fl*^ *Vil1*^*Cre-*^) or deficient *(Il6ra*^*fl/fl*^ *Vil1*^*Cre+*^). Confocal imaging confirmed selective ablation of IL-6R*α* in IECs in *Il6ra*^*fl/fl*^ *Vil1*^*Cre+*^ mice (**fig. S4, A and B**). Following injection of IL-6 during pregnancy, intestinal Th17 cells were significantly reduced in the gut of *Il6ra*^*fl/fl*^ *Vil1*^*Cre+*^ offspring, compared to *Il6ra*^*fl/fl*^ *Vil1*^*Cre-*^ littermate controls (**Fig. 4F**). Together, these results demonstrate that IL-6 signaling in fetal IECs is sufficient and necessary to confer long-term intestinal immune alterations.

Prior lineage tracing studies revealed that all fetal IECs can contribute to the adult intestinal epithelial stem cell (EpSC) pool that is characterized by *Lgr5* expression *(33)*. To address the possibility that alterations in fetal IECs may have long-term consequences on adult EpSCs, we assessed the chromatin accessibility of EpSCs (EpCAM^+^ Lgr5^+^) from 6-week-old offspring delivered by control or IL-6-injected dams using ATAC-seq (**Fig. 5A and fig. S5A**). Chromatin accessibility was increased in EpSCs of adult offspring delivered by IL-6-injected dams, with 1006 regions uniquely open (fold change greater than 2), compared to only 71 regions in EpSCs of control offspring (**Fig. 5B**). As anticipated based on the profound changes in epithelial cells occurring post-birth and during weaning *(34, 35)*, open chromatin sites were largely distinct from those observed in the fetuses, with only 17 common peaks (< 1% of the fetus) shared between fetal IECs and adult EpSC (**fig. S5B**). Differentially accessible promoters found in adult EpSCs of offspring from IL-6-injected dams showed enrichment of genes involved in MAPK activation, cytokine signaling and defense responses (e.g. *Alox5ap, Ptgs2, Mgst2, Ffar3)* (**Fig. 5C**).

**Figure 5.**
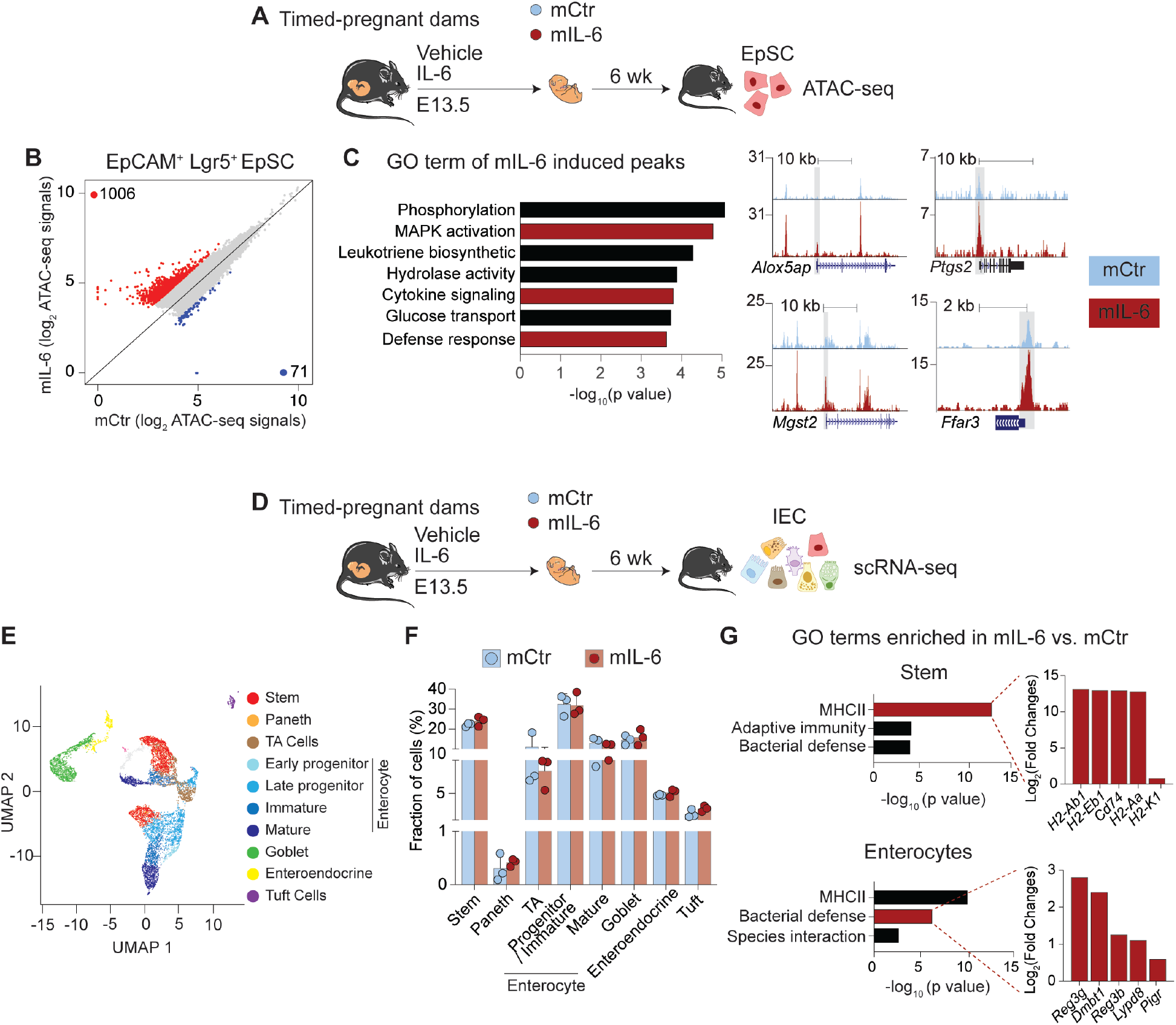
Increasing maternal IL-6 levels during pregnancy alters the chromatin accessibility and transcriptome of offspring intestinal epithelial stem cells. (**A-C**) (**A**) Timed-pregnant dams were injected with vehicle or 10 μg of IL-6 at E13.5. Intestinal epithelial stem cells (EpSC) were sorted from their offspring, mCtr or mIL-6, for ATAC-seq at 6 weeks old. (**B**) Scatter plot showing open chromatin regions. Red and blue dots correspond to regions that were differentially accessible by >2 fold in the EpSCs of mIL-6 and mCtr offspring, respectively. (**C**) Left: Gene Ontology (GO) term analysis of differentially accessible promoter regions (peaks within 3kb of the next transcription start site, fold change > 2) in mIL-6 EpSC compared to mCtr EpSC. Right: UCSC Genome Browser snapshots of genes involved in cytokines signaling. Promoters open in mIL-6 offspring are highlighted in gray boxes. (**D-G**) (**D**) Intestinal epithelial cells (IEC) were sorted from mCtr or mIL-6 offspring for scRNA-seq at 6 weeks old. (**E**) UMAP projection of IEC subsets, assigned based on the 25 most differentially expressed genes in each subset versus all other cells. (**F**) Frequency of each IEC subset. (**G**) Top 3 GO terms that were enriched in stem cells and enterocytes of mIL-6 offspring. For selected GO terms highlighted in red, the top 5 genes upregulated in mIL-6 offspring are shown. Data are from 1 experiment with 3 pregnant dams per group, using 3-5 offspring per pregnancy. See also Figure S5 and S6.

In adults, intestinal EpSCs constantly self-renew and differentiate into diverse subsets of epithelial cells endowed with discrete functions *(33)*. We next evaluated the potential consequence of maternal-IL-6-mediated EpSC alterations on the composition and transcriptome of IECs. We focused our analysis on the gut compartments with the largest increase in Th17 cells post-maternal IL-6 injection (duodenum and ileum) (**Fig. 3D and 5D**). IECs isolated from offspring delivered by control (n=3, 4929 cells) and IL-6-injected (n=3, 4112 cells) dams were evaluated via droplet-based 3’ single cell RNA-sequencing (scRNA-seq). Unsupervised clustering partitioned the cells into 21 clusters. Each cluster was identified (**Fig. 5E, fig. S6, A and B)** based on signature genes associated with distinct cell types and differentiation states *(36)*. No significant differences were observed in cell composition between offspring delivered by control and IL-6-injected dams (**Fig. 5F**). On the other hand, IL-6 injection imposed discrete transcriptomic alterations in defined subsets of IECs, including stem cells, transit-amplifying (TA) cells and subsets of enterocytes at various stages of differentiation (**fig. S6C**). In particular, genes encoding canonical components of the MHCII machinery, including *Cd74, H2-Aa* and *H2-Ab1* and, were strongly enriched in both stem cells and enterocytes of offspring delivered by IL-6-injected dams compared to control dams, and to a lesser extent in the ileal enteroendocrine and goblet cells (**Fig. 5G and fig. S6C**). Of note, expression of MHCII and genes encoding for the antigen presentation machinery by EpSCs have recently been reported to be microbiota-dependent and to contribute to epithelial cell differentiation *(37, 38)*. IL-6 injection during pregnancy was also associated with increased expression of genes encoding antimicrobial peptides *(Reg3b, Reg3g)* in enterocytes (**Fig. 5G**). Thus, an increase in IL-6 during pregnancy can have a long-term impact on EpSC chromatin accessibility and epithelial cell function.

We next determined the physiological impact of our observations on host fitness. Altered epithelial activation status and enhanced T cell responses to the microbiota in the gut of offspring of previously infected or IL-6-injected dams pointed to the possibility that this phenomenon may be associated with enhanced antimicrobial defense. To address this hypothesis, we employed an acute model of oral infection with *Salmonella enterica* serovar Typhimurium *(S*. Typhimurium) (**Fig. 6A**). At the dose and route of infection used, *S*. Typhimurium primarily invades intestinal epithelial cells before disseminating to distal organs and causing host lethality *(39)*. While no differences in bacterial shedding were observed at days 1 and 2 post-infection, by day 4, offspring of IL-6-injected dams displayed a 1 to 2-log reduction in bacterial burden in their feces and Peyer’s patches compared to the offspring of control dams (**Fig. 6, B and C**). Quantification of fecal lipocalin-2 revealed that infection-induced pathology was also significantly reduced in the offspring of IL-6-injected dams compared to control dams (**Fig. 6D**). While the direct role of IL-17A in controlling *S*. Typhimurium remains unclear *(40)*, infection with this bacterium has been associated with early Th17 responses *(41)*. At day 5 post-infection, the number and frequency of IL-17A-and IL-17A/IL-22-producing CD4^+^ T cells, but not of ILCs, were significantly increased in the siLP of offspring of IL-6-injected dams compared to control dams (**Fig. 6E, fig. S7A**). Enhanced IL-6 during pregnancy also reduced bacterial dissemination to distal organs in offspring, with a 2 to 3-log decrease in bacteria burden in the mesenteric lymph nodes, the liver and the spleen compared to controls (**Fig. 6F**). Finally, injection of IL-6 to dams significantly delayed lethality and improved offspring survival rates post-infection (**Fig. 6G**). Together, our data support the idea that enhanced IL-6 signaling experienced by the fetus during pregnancy can promote antimicrobial defense via its impact on epithelial responses and/or T cell responses.

**Figure 6.**
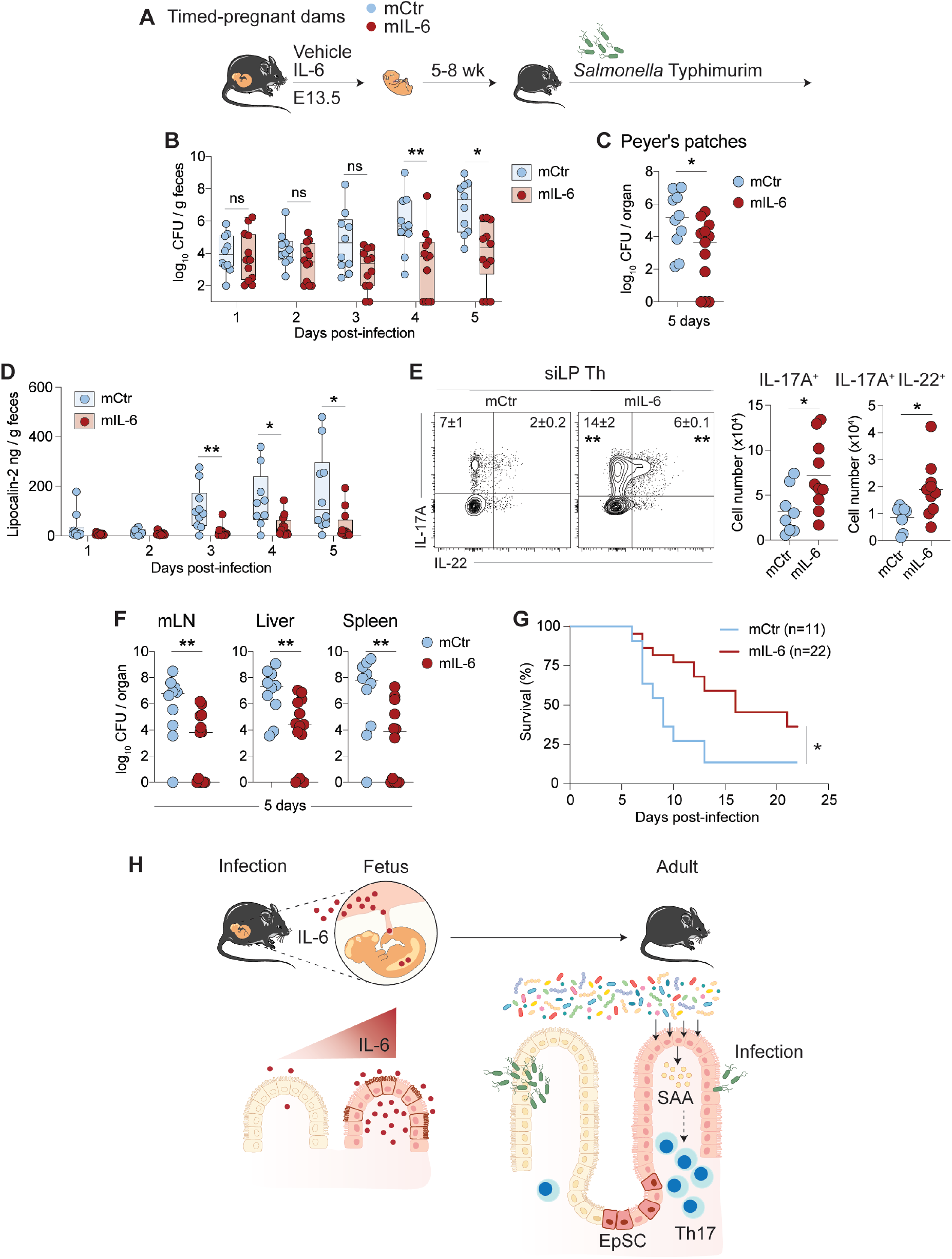
Increasing maternal IL-6 levels during pregnancy enhances protection to gastrointestinal infection in offspring. (**A**) Timed-pregnant dams were injected with vehicle or 10 μg of IL-6 at gestation day 13.5 (E13.5), and their offspring (mCtr or mIL-6, respectively) were orally administered with *Salmonella* Typhimurim *(S*. Typhimurim) at 5 to 8 weeks old. (**B-C**) *S*. Typhimurium burden in (**B**) feces at days 1 to 5 post-infection and (**C**) Peyer’s patches at day 5 post-infection. (**D**) Fecal lipocalin-2 at days 1 to 5 post-infection. (**E**) Left: Representative contour plots showing cytokine production by siLP Th (TCRβ^+^ CD4^+^ Foxp3^-^). Right: Total IL-17A^+^ and IL-17^+^ IL-22^+^ siLP Th cells. (**F**) *S*. Typhimurim burden in in distal tissues at day 5 post-infection. (**G**) Survival curve. (**H**) Model: direct responses of fetal intestinal epithelial cells to IL-6 during maternal infection confer an enduring epigenetic memory on adult intestinal stem cells. As a result, offspring epithelial cells express higher levels of antimicrobial peptides and are able to promote and/or sustain heightened antimicrobial adaptive response for the long-term. (**A-H**) The CFU detection limit was 10^2^ CFU per gram or organ. Data are representative of 3 independent experiments with 1-2 pregnant dams per group, using 3-6 offspring per pregnancy. Numbers in representative flow plots indicate mean ± SD. Each dot represents an individual mouse. * p < 0.05; ** p < 0.01; ns, not significant (two-tailed unpaired Student’s t test in B-F or a Log-rank test in G). See also Figure S7.

Optimal responses to microbial challenges are of utmost importance for host survival. Here we showed that pregnancy represents a critical stage for tissue-specific immune education of the offspring. Notably, we demonstrated that direct responses of fetal epithelial cells to cytokines produced during maternal infections confer an enduring epigenetic memory to gut stem cells. As a result, epithelial cells express higher level of antimicrobial peptides and are able to promote and sustain heightened antimicrobial adaptive T cell responses (**Fig. 6H**). Recent studies have uncovered that, in adults, epithelial stem cells can develop memory of inflammatory insults, a phenomenon associated with accelerated responses to subsequent injuries *(2, 3, 42)*. Our present work proposes that even a transient, mild infection encountered during prenatal development can impose lasting alterations to gut epithelial stem cells, resulting in heightened protective immunity.

Of particular interest, the impact of maternal infection was tissue-specific and dominantly mediated by a single cytokine, IL-6. This observation raises an intriguing possibility that during fetal development, stem cells residing in different compartments may be educated by highly specific signals to develop optimal tissue immunity. Our findings also support the idea that infections experienced by the mother during pregnancy may optimize responses to immediate microbial threats in the offspring by augmenting responses at the target infectious sites. Of note, IL-6 has previously been reported to be increased throughout pregnancy *(43)*, a phenomenon that may broadly contribute to the education of the mammalian gut immune system.

While our data indicate that consequences of maternal microbial exposure can be coopted by the fetus to develop optimal immune fitness, altered thresholds of immune activation in tissues may also, in some settings, promote the development of inflammatory responses. Further explorations of the maternal-fetal dialogue in the face of microbial or environmental challenges may provide new insights into our understanding of homeostatic and protective tissue immunity, as well as host predispositions to inflammatory diseases.

## Acknowledgements

We thank the NIAID animal facility, the NIAID Microbiome Program gnotobiotic animal facility, Teresa Hawley, Jinguo Chen, Shreni Mistry, Galina Koroleva, Kimberly Beacht and Ejae Lewis for technical support. We thank Kairui Mao, Anita Gola and the NIAID Biological Imaging Facility for providing support for confocal imaging. We thank all Belkaid laboratory members, especially Samira Tamoutounour, Pete Kulalert, Michel Enamorado and Eduard Ansaldo for providing constructive feedback to this project and critical reading of this manuscript.

## Author contributions

AIL and YB designed the study experiments and wrote the manuscript. AIL performed experiments and analyzed the data. TM and TF participated in performing experiments. VL analyzed RNA-seq and ATAC-seq data. AS provided expertise in *Salmonella* Typhimurium infection experiment and analyzed 16S data. HYS provided expertise in ATAC-seq. OJH provided intellectual expertise. RMK and HAC performed and interpreted data from behavior studies. SJH provided intellectual expertise, participated in experiments and helped to interpret experimental results.

## Data and materials availability

16S sequencing data, ATAC-seq, RNA-seq and scRNA-seq data were deposited into the NCBI Sequence Read Archive (accession number PRJNA666968, GSE158950, GSE159149).

## Supplementary Materials

## Materials and Methods

### Mice

Specific Pathogen Free (SPF) C57BL/6, *Lgr5*^*EGFP-IRES-creERT2*^ (B6.129P2-*Lgr5*^tm1(cre/ERT2)/Cle^/J, Jax008875), *Il6ra*^fl/fl^ (B6.SJL-*Il6ra*^tm1.1Drew^/J, Jax012944), *Vil1*^cre^ (B6.Cg-Tg(Vil1-cre)1000Gum/J, Jax021504) mice were purchased from The Jackson Laboratory and Taconic Biosciences (SFB negative barrier). Germ-free C57BL/6 mice were bred and maintained in the NIAID Microbiome Program gnotobiotic animal facility. CBir1 Tg mice were generated by Dr. Charles Elson (University of Alabama, Birmingham, AL), obtained under a material transfer agreement and back-crossed to CD45.1-expressing mice *(1)*. All mice were bred and maintained at an American Association for the Accreditation of Laboratory Animal Care (AAALAC)-accredited animal facility at NIAID and housed in accordance with procedures outlined in the Guide for the Care and Use of Laboratory Animals. All experiments were performed at NIAID under an animal study proposal (LISB19E) approved by the NIAID Animal Care and Use Committee. For timed-pregnant breeding, males were first singly housed for 3 days and cohoused with females for breeding. Males were separated from the breeding cage once a plug was observed. Littermate dams were used in each experiment to help normalize for differences in the microbiota. Offspring were weaned at 3 weeks after birth. Age-matched mice between 5 to 8 weeks from both sexes were used in each experiment unless specified otherwise.

### *Yersinia pseudotuberculosis* infection

The *ΔyopM* mutant strain of *Yersinia pseudotuberculosis* 32777 was grown overnight in 2X YT media (Sigma) at 25°C with shaking at 200 rpm. Overnight cultures were centrifuged at 3000 g, resuspended in PBS and adjusted to a density (absorbance at 600nm (A_600_)) of 0.1 in PBS. Mice fasted overnight (12-16 hours) in new cages were orally administered 200 μl of *yopM* suspension (∼2 × 10^7^ colony forming unit (CFU)). To determine bacterial burden in the maternal spleen, liver, placenta and fetal liver after *yopM* infection, tissues were homogenized through 70 μm filters with PBS. Tissue homogenates were serially diluted and plated onto MacConkey plates, and colonies were counted after incubation at 25°C for 48h. To measure bacterial burden in feces, fecal DNA was extracted by phenol/chloroform, and the *yscF* gene was quantified against a standard curve by qPCR (forward primer 5’-ATGAGTAACTTCTCTGGATTTACG-3’, reverse primer 5’-TTATGGGAACTTCTGTAGGATG-3’).

### *Salmonella* Typhimurium infection

A nalidixic acid-resistant strain of *S*. Typhimurium, IR715 (provided by Prof. Andreas Bäumler), was cultured in LB at 37°C with shaking at 200 rpm. Overnight cultures were centrifuged at 3000 g, resuspended in PBS and adjusted to an A_600_ of 0.001. Mice were fasted overnight (12-16 hours) prior to oral administration of 200 μl of *S*. Typhimurium suspension (∼2 × 10^5^ CFU). Fecal samples were collected into pre-weighed tubes, homogenized in 500 μl of PBS with pipette tips and vortexed for 5 seconds. At day 5 post-infection, collected tissues were homogenized through 70 μm filters with PBS. The top fraction of fecal and tissues homogenates was serially diluted in PBS, spotted with a multichannel pipette onto LB agar with 50 μg/ml nalidixic acid and incubated overnight at 37°C prior to counting CFU. Remaining fecal homogenates were centrifuged at 3000 g and stored at -80°C for lipocalin-2 measurement.

### Adoptive transfer of CBir1 Tg CD4^+^ T cells

Cells from the spleen and the lymph nodes of CD45.1 CBir1 Tg mice were isolated by being passed through 70 μm filters. To enrich for CD4^+^ T cells, single-cell suspensions were incubated with biotin-conjugated anti-CD8, anti-NK1.1, anti-CD11b and anti-CD19 antibodies followed by anti-biotin microbeads (Miltenyi) and negatively selected by magnetic cell separation with MACS technology (Miltenyi). 0.5 – 1.0 × 10^6^ enriched CD4^+^ T cells from CBir1 Tg mice were intravenously transferred via retroorbital injection to offspring delivered by control or transiently infected dams on a congenic CD45.2 background. Recipient mice were sacrificed at day 10 to 12 post transfer to assess donor cells in the small intestinal lamina propria.

### Cross-fostering experiments

Within 24 hours after birth, whole litters were removed from the original mother and gently mixed with new bedding in clean cages. Fosters mothers were held to urinate on the pups. The cages were covered to avoid disturbances and monitored for at least 48 hours. Pups were nurtured by foster mothers until weaning at 3 weeks after birth.

### Serum transfer experiments

Whole blood was collected from *yopM-*infected timed-pregnant dams at days 3 and 5 post-infection. Blood samples were incubated at room temperature for 30 min and centrifuge at 2000 g for 10 minutes. Sera were collected and transferred immediately via retroorbital injection to naïve timed-pregnant dams at gestation day 13.5 and 15.5. Each recipient received 200 μl of sera, combined from at least two mice, at each time point. 50 μl of serum collected from each mouse was plated onto MacConkey plates to assess CFU.

### *In vivo* cytokine stimulation and blockade

Timed-pregnant mice dams were intravenously injected with up to 10 μg of recombinant mouse cytokines (Biolegend) at gestation day 13.5. For IL-6 blockade experiments, timed-pregnant dams at gestation day 10.5 were injected intraperitoneally with either 1 mg of anti-IL-6 (MP5-20F3, BioXCell) or the IgG1 isotype control (HPRN, BioXCell) 1 hour before *yopM* infection.

### Germ-free mice conventionalization

Offspring delivered by PBS or IL-6-injected germ-free timed-pregnant mice were kept in germ-free isolators and fostered by the same mothers until weaning. Subsequently, half of the litter was kept in isolators, while the other half was transferred using aseptic technique to a SPF facility at 3 weeks old and analyzed at 5-8 weeks old.

### Tissue processing

Mice were euthanized with CO_2_, perfused via the left ventricle of the heart with 10 ml of PBS, and the siLP, cLP, lung and skin were collected and placed into cold complete medium (RPMI 1640 supplemented with 2 mM L-glutamine, 1 mM sodium pyruvate, 1 mM nonessential amino acids, 20 mM HEPES, 50 mM β-mercaptoethanol, 100 U/mL penicillin, 100 mg/mL streptomycin). For siLP and cLP preparation, the Peyer’s patches and the mesenteric adipose tissue were removed. Tissues were opened, washed in cold PBS to remove feces, cut into 1-2 cm segments and treated with complete medium containing 5 mM EDTA and 0.145 mg/ml dithiothreitol for 20 min at 37°C with constant stirring. Tissues were shaken vigorously for 1 min and further digested with 10 ml of medium containing 500 μg/ml DNase I (Sigma-Aldrich) and 100 μg/ml Liberase TL (Roche) with continuous stirring at 37°C. Lungs were diced and incubated in 2 ml of pre-warmed medium containing 1000 μg/ml DNase I and 250 μg/ml of Liberase TL for 25 min at 37°C. To isolate cells from the skin, ear pinnae were excised and separated into ventral and dorsal sheets, digested by placing the dermal side down in media containing 500 μg/ml DNase I with 250 μg/ml Liberase TL and incubated for 90 min at 37°C. Digested intestine, lung and skin were passed through 70 μm cell strainers. Leukocytes were enriched by resuspension in 4 ml 37.5% Percoll and centrifuged at 1800 rpm for 5 min. Cells were then washed with PBS before downstream analysis.

### *In vitro* re-stimulation

To assess cytokine production potential, single-cell suspensions were restimulated in complete medium containing 10% FBS, 50 ng/mL phorbol myristate acetate (PMA; Sigma-Aldrich), 5 mg/mL ionomycin (Sigma-Aldrich) and a 1:1000 dilution of GolgiPlug (BD Bioscience) for 2.5 h at 37°C.

### Flow cytometry analysis

Fluorophore-conjugated antibodies that were used are listed in Table S1. For intracellular cytokine and transcription factor staining, after surface staining, cells were fixed and permeabilized using the Foxp3/Transcription Factor Staining Buffer Set (ThermoFisher) for at least 1 hour at 4°C and stained with fluorophore-conjugated antibodies for at least 1 hour at 4°C. All staining was performed in the presence of purified anti-mouse CD16/32 and purified rat gamma globulin. Dead cells were excluded using 4’,6-diamidino-2-phenylindol (DAPI, Sigma-Aldrich) or LIVE/DEAD Fixable Blue Dead Cell Stain Kit (Invitrogen Life Technologies). For phospho-STAT3 staining, after surface staining, cells were fixed with 4% paraformaldehyde at room temperature for 10 minutes and permeabilized in pre-chilled methanol at -80°C for 1 hour. Cells were subsequently washed with PBS containing 0.5% Triton X-100, 0.1% BSA and stained with a PE-conjugated phospho-STAT3 antibody (Cell Signaling) or the isotype control in the same buffer at 4°C. Cells were washed and analyzed immediately. Integrated mean fluorescence intensity (iMFI) reflects the total functional responses of a population that determined by amount of expression level (MFI) and frequency of positive cells *(2)*.

### 16S rRNA gene profiling

Fecal DNA was extracted using MagAttract PowerMicrobiome DNA/RNA kit (Qiagen). 16S sequencing libraries were generated using 100 ng of purified DNA as template and the V4 region-targeting primers 515F (5′-GTGCCAGCMGCCGCGGTAA-3′) and 806R (5′-GGACTACHVGGGTWTCTAAT-3′). Amplicons were quantified, pooled at equimolar concentrations and sequenced on a MiSeq (Illumina) using the v3 MiSeq Reagent Kit (Illumina). 16S data were analyzed using QIIME 2 *(3)*. Briefly, reads were denoised using the q2-dada2 plugin (exact parameters were forward reads truncated to 180 bases with 20 bases trimmed from the 5’ end, reverse reads truncated to 150 bases with 20 bases trimmed from the 5’ end, and bases truncated with quality score 10). Using the q2-feature-table plugin, samples within an experiment were rarefied to the depth of the sample with the fewest reads, and sequence variants were filtered if within an experiment they did not occur in at least 4 samples. Rooted phylogenetic trees were generated using the align-to-tree-mafft-fasttree method in the q2-phylogeny plugin. Alpha and beta diversity analyses were performed using the core-metrics-phylogenetic method in the q2-diveristy plugin. Taxonomy was assigned to sequence variants using the q2-feature-classifier plugin and a naïve Bayes classifier pre-trained on the Silva 132 515F/806R 99% OTUs provided as a QIIME 2 data resource. A pseudocount was applied prior to calculating differences in the relative abundance of taxonomic groups.

### Confocal microscopy

Fetal intestines were isolated using a dissecting microscope. Adult small intestines were flushed with cold PBS and prepared using the Swiss roll technique. Samples were incubated overnight in fixation and permeabilization buffer (BD Bioscience) followed by dehydration in 30% sucrose before embedding in OCT compound (Sakura Finetek). 20-μm sections were cut on a CM3050S cryostat (Leica) and adhered to Superfrost Plus slides (VWR). Frozen sections were treated with pre-chilled acetone for 30 min at -80°C, subsequently permeabilized and blocked in PBS containing 0.3% Triton X-100 and 1% BSA (Sigma-Aldrich). Samples were stained with anti-IL6R*α* (BD) and anti-EPCAM (Biolegend) at 37°C for 30 min. Images were captured on a Leica TCS SP8 confocal microscope equipped with HyD and PMT detectors and a 40X oil objective. Images were analyzed using Imaris (Bitplane).

### Intestinal epithelial cells isolation and sorting

Freshly isolated and longitudinally opened samples were washed twice with gut wash buffer (HBSS supplemented with 2 mM L-glutamine, 1 mM sodium pyruvate, 1 mM nonessential amino acids, 20 mM HEPES, 100 U/mL penicillin, 100 mg/mL streptomycin and 2% FCS). Tissues were cut into 1-2 cm segments and incubated with gut wash buffer containing 2 mM DTT in a 37°C water bath for 10 minutes. Tissues were vigorously shaken and subsequently incubated with fresh gut wash buffer containing 5 mM EDTA in a 37°C water bath for 7 minutes. EDTA-containing fractions were collected, and tissues were incubated with fresh EDTA for another 7 minutes. The 2 EDTA-containing fractions (enriched with crypts) were passed through 100-μm filters and centrifuged at 300 g for 5 min. Cell pellets were digested with DMEM supplemented with 0.1 U/ml Dispase (Life Technology) and 100 μg/ml DNase I for 8 minutes in a 37°C water bath. Digestion was neutralized with 2.5% FCS-DMEM, and the pellets were washed twice with gut wash buffer. Cell pellets were stained with antibodies against EpCAM, lineage (TCRβ, TCRγδ, CD8, CD11b), CD31 and DAPI on ice for 30 minutes. For single-cell RNA-sequencing, TotalSeq-A hashtag oligonucleotides (Biolegend) were included along with surface antibodies. Intestinal epithelial cells were sorted as the DAPI^-^ CD31^-^ Lineage^-^ EpCAM^+^ population using a MA900 cell sorter (Sony) with a 100-μm chip.

### Whole-tissue RNA-seq

3 cm of ileum was collected and extensively washed with PBS to remove feces. Samples were homogenized by bead-beating in tubes containing 2.8mm beads and TRIzol (Thermofisher). RNA was extracted according to the TRIzol protocol. cDNA was synthesized with the Ultralow V4 kit (Clonetech), and sequencing libraries were subsequently prepared with the Nextera XT DNA prep kit (Illumina). RNA-seq reads were mapped with STAR to the mouse reference genome (mm10) with default parameters. Gene expression was measured with HOMER using analyzeRepeats.pl with parameters -condenseGenes and -tpm (for scatter plots) or -noadj (for differential gene expression analysis). Differential gene expression was assessed with HOMER’s getDiffExpression.pl utilizing DESeq2. Genes with a fold change greater 2 and FDR smaller than 0.05 were considered differentially expressed.

### FAST-ATAC-seq

Fast-ATAC-seq was performed according to a published protocol *(4)*. Briefly, 10,000 fetus IEC (EpCAM^+^) or adult EpSC (EpCAM^+^ Lgr5^+^) were sorted and pelleted by centrifugation at 500 g for 5 min at 4°C. Supernatant was removed, and cell pellets were resuspended in 45 μl of transposase mixture (25 μl Tagment DNA Buffer, 2.5 μl Tagment DNA enzyme, 0.5 μl 1% digitonin and 17 μl H_2_O) and incubated for 30 minutes at 37°C with agitation at 300 rpm. Tagmented DNA was purified using the QIAGEN MinElute Reaction Cleanup Kit, and purified DNA was eluted in 10 μl of elution buffer. Tagmented fragments were amplified with 10-15 PCR cycles based on an amplification curve. After purification using the QIAGEN PCR cleanup kit, samples were sequenced single-end on a HiSeq 2500. ATAC-seq reads from 3 biological replicates were demultiplexed using bcl2fastq and mapped to the mouse reference genome (mm10). All biological replicates were run through the Irreproducible Discovery Rate (IDR) framework, and only replicates that passed all filters were kept. Peaks that passed IDR filter criteria (<0.05) were kept. For final analysis, all peaks that passed IDR filter and were found in all replicates were used. In total there were 24938 peaks in adult EpSC cells from mCtr samples and 30614 peaks in adult EpSC cells from mIL-6 samples. For analysis of the fetal IECs 49787 peaks were found in mCtr samples and 43813 in mIL-6 samples. For the IDR pipeline, open regions of chromatin were called using HOMER *(5)*.

### Single-cell RNA sequencing and transcriptome analysis

2.5 × 10^4^ of EpCAM^+^ intestinal epithelial cells were isolated by a Sony MA900 cell sorter from each sample. All samples were labelled with TotalSeqA hashtags antibodies (Biolegend), pooled and encapsulated into droplets using a Chromium Single Cell Controller (10X Genomics), and libraries were prepared using Chromium Single Cell 3 Reagent Kits v3 (10X Genomics). The Controller was loaded with 2.5 × 10^4^ cells per lane. mRNA was prepared following the 10X Genomics user guide, while the HTO library was prepared according to published guidelines *(6)*. Libraries were sequenced on a NextSeq 500 with 10% of the lane occupied by the HTO library and 90% by the mRNA library. HTO was assigned to each single cell. Data were normalized, and lanes were combined using the scTransform function in Seurat. Data were displayed as uniform manifold approximation and projection (UMAP) using 25 dimensions. Intestinal cell subsets from each cluster were defined by the top 25 differentially expressed genes and identified based on published datasets *(7)*. Gene expression within each cluster was compared between offspring delivered by control or IL-6-exposed dams using Seurat.

### Three-chamber social behavior tests

8-week-old offspring were tested for sociability by three-chamber social tests. Mice were habituated in the center chamber for 10 min, followed by a 10 min habituation session with access to all three empty chambers. After the two habituation sessions, the 10 min sociability test was conducted. An inverted wire pencil cup (novel object) was placed in one of the side chambers and the other side chamber had an inverted wire pencil cup containing an age- and sexed-matched novel C57Bl6/J mouse (novel mouse). Frequency to enter and time in each chamber was analyzed using Noldus Ethovision software (Noldus Information Tech Inc. Leesburg, VA, USA).

### Measurement of cytokines and chemokines

Sera cytokine and chemokine levels were quantified by the Cytokine & Chemokine Convenience 36-Plex Mouse ProcartaPlex Panel 1A (Invitrogen) according to the manufacturer’s instructions.

### Measurement of serum amyloid A

3 cm of intestinal segments from the duodenum, jejunum and ileum were collected and extensively washed with PBS to remove feces. Samples were homogenized by bead-beating with 400 μl of RIPA lysis buffer (Chem Cruz) containing proteinase inhibitor in tubes containing 2.8mm beads. Extracted samples were centrifuged at 10,000 g for 10 min at 4°C to remove debris. Serum Amyloid A levels were quantified by the Mouse Serum Amyloid A Quantikine ELISA Kit (R&D) according to the manufacturer’s instructions.

### Measurement of lipocalin

Lipocalin-2 levels in fecal samples were measured by the Mouse Lipocalin-2/NGAL Quantikine ELISA Kit (R&D) according to the manufacturer’s instructions.

### Statistical analysis

Groups were compared with Prism software (GraphPad).

---

**Figure S1.**
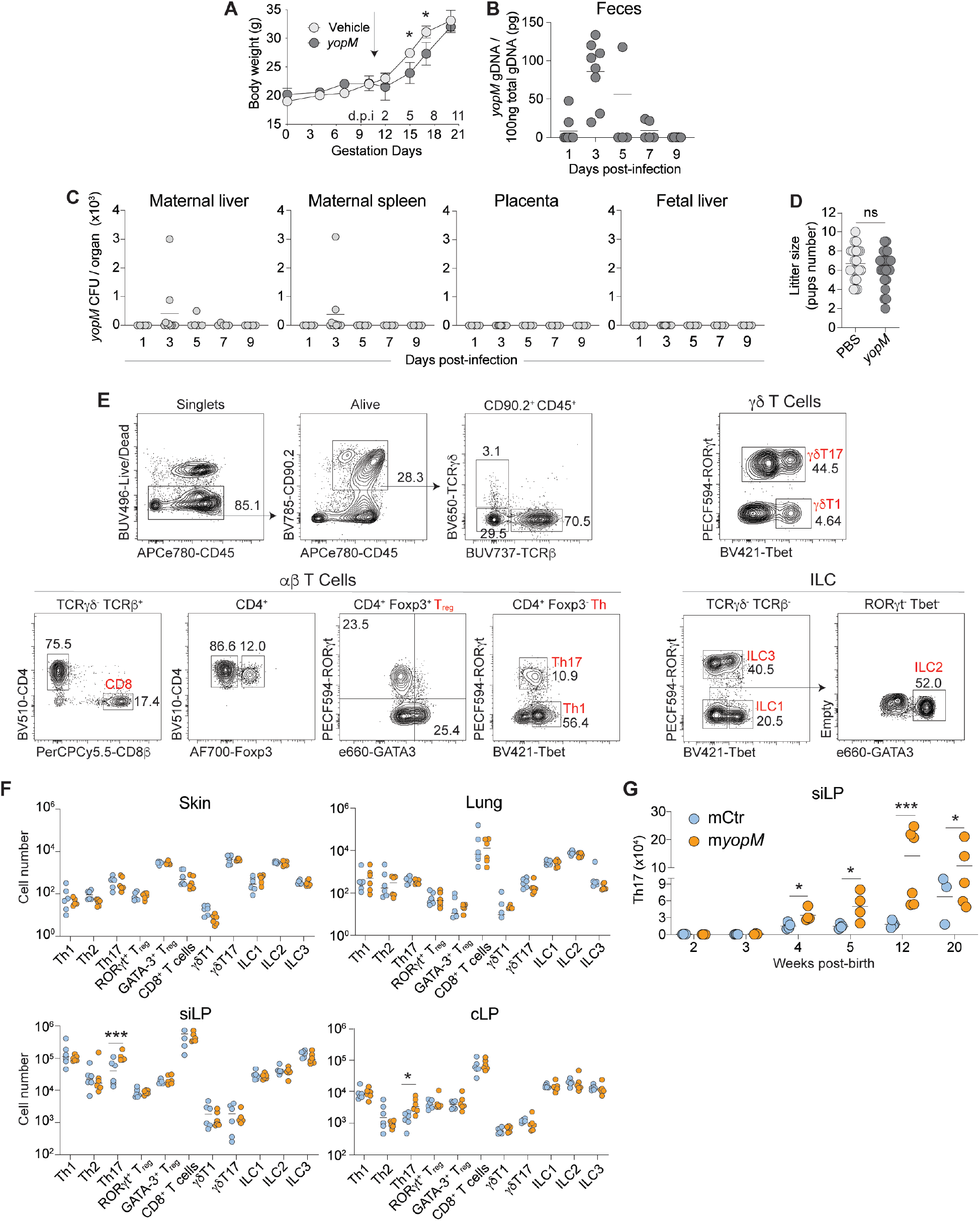
Maternally restricted infection induces tissue-specific alterations to offspring immunity. (**A-D**) Timed-pregnant dams were orally administered with vehicle (PBS) or *Y. pseudotuberculosis ΔyopM (yopM)* at gestation day 10.5 (E10.5). (**A**) Bodyweight of vehicle-treated or *yopM*-infected dams were measured over the course of pregnancy. (**B**) *yopM* burden in feces at days 1 to 9 post-infection. (**C**) *yopM* burden in maternal tissues, placenta and fetal liver at days 1 to 9 post-infection. (**D**) Number of pups per litter from vehicle-treated or *yopM*-infected dams. (**E**) Flow cytometry plots depict the gating scheme of lymphocyte subsets in small intestinal lamina propria (siLP) and applicable to other organs including colonic lamina propria (cLP), ear skin and lung. (**F**) Absolute cell number of lymphocyte subsets in skin, lung, siLP and cLP of offspring from vehicle-treated dams (mCtr) or *yopM*-infected dams *(myopM)*. (**G**) Total siLP Th17 cell number in the mCtr or m*yopM* offspring at corresponding ages. (**A-F**) Data combines 3 independent experiments (A-C), each experiment with 1-3 pregnant dams per group, or representative of 3 independent experiments (E-G) with 1-2 pregnant dams per group, using 3-5 offspring per pregnancy, or pool from all experiments (D). Each dot represents an individual mouse. Numbers in representative flow plots indicate mean. * p < 0.05; *** p < 0.001; ns, not significant (two-tailed unpaired Student’s t test).

**Figure S2.**
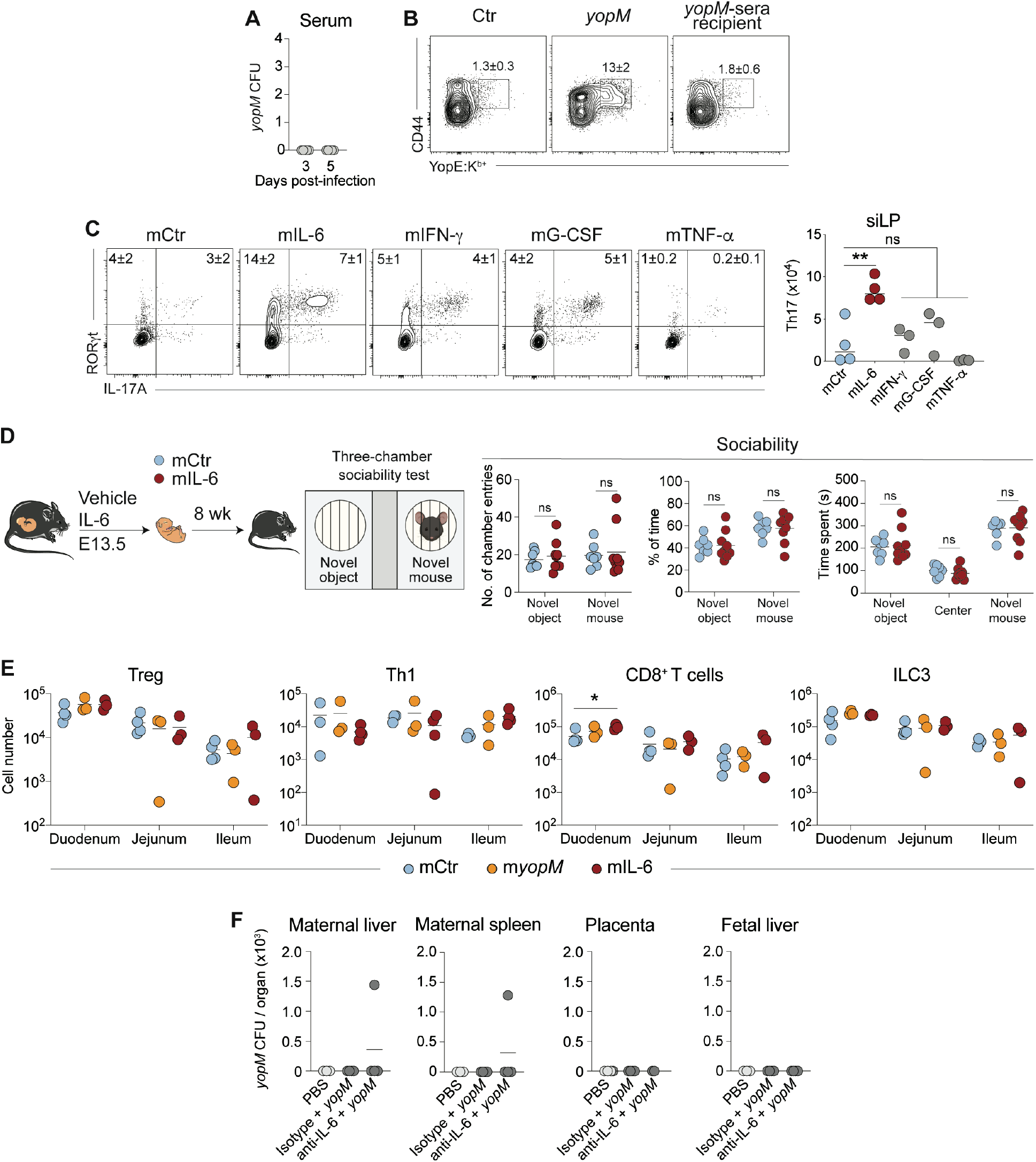
Increasing maternal IL-6 levels during pregnancy is sufficient to enhance offspring intestinal Th17 and responsiveness to the microbiota. (**A**) *yopM* burden in pregnant dams’ sera at days 3 and 5 post-infection. (**B**) Representative contour plots showing CD44 and YopE:K^b+^ expression by splenic CD8^+^ T cells of vehicle-treated, infected or *yopM*-sera recipient dams at 4-weeks post-treatment. (**C**) Timed-pregnant dams were intravenously injected with vehicle or 2 μg of indicated cytokines at gestation 13.5, and their offspring were analyzed at 6 weeks old for transcription factor expression and cytokine production after restimulation. Left: Contour plots were gated on live CD45^+^ CD90.2^+^ TCRβ^+^ CD4^+^ Foxp3^-^. Right: Total Th17 cell number in the siLP. (**D**) Timed-pregnant dams were injected with vehicle or 10 μg IL-6 injected at E13.5, and their offspring, mCtr or mIL-6, were subjected to three-chamber sociability tests. Sociability was assessed by number of times enter each chamber and time spent at each chamber. (**E**) siLP sub-compartments from 6-week-old mCtr, m*yopM* offspring, or the offspring of dams IV injected with 10 μg of IL-6 (mIL-6) were analyzed for total number of lymphocyte subsets. (**F**) *yopM*-infected dams were pre-treated with 1 mg of anti-IL-6 monoclonal antibody or the isotype control 2 hours prior to infection. *yopM* burden in maternal tissues, placenta and fetal liver at days 3 post-infection. (**A-F**) Data are representative of 3 independent experiments (C, E), each experiment with 1-2 pregnant dams per group, using 3-4 offspring per pregnancy, or 1 experiment (D) with 1-2 pregnant dams per group using 3-7 offspring per pregnancy, or pool 3 independent experiments (A, B, F), each experiment with 1-3 pregnant dams per group. Each dot represents an individual mouse. Numbers in representative flow plots indicate mean ± SD. * p < 0.05; ** p < 0.01; ns, not significant (ANOVA multiple-comparison test, one-way in C, F and two-way in D, E).

**Figure S3.**
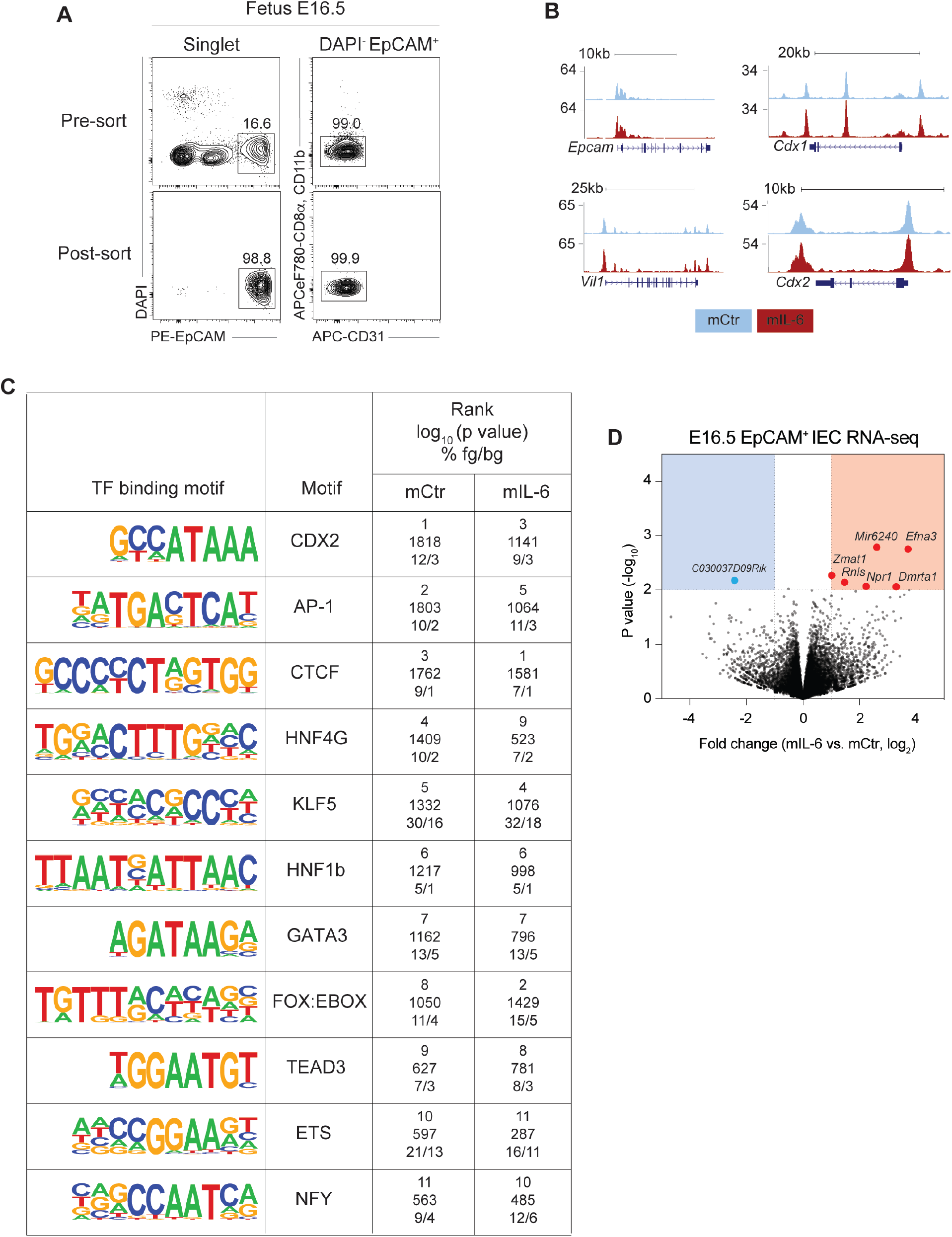
Increasing maternal IL-6 levels during pregnancy alters the epigenome of fetal intestinal epithelial cell. (**A**) Flow plots showing strategy to isolate fetal intestinal epithelial cells (IEC, EpCAM^+^) and post-sort purity check. (**B**) UCSC Genome Browser snapshots of IEC signature genes. (**C**) Enrichment analysis for transcription factor binding motifs of fetal IEC. (**D**) Volcano plot of fetal IEC transcriptomic profiles comparing fetal IEC from IL-6-treated dams versus control dams. Upregulated genes from fetuses IEC of IL-6 injected dams are denoted in red, downregulated gene is denoted in blue. Data are from 1 experiment (D-E) with 3 pregnant dams per group, using 4-7 offspring per pregnancy.

**Figure S4.**
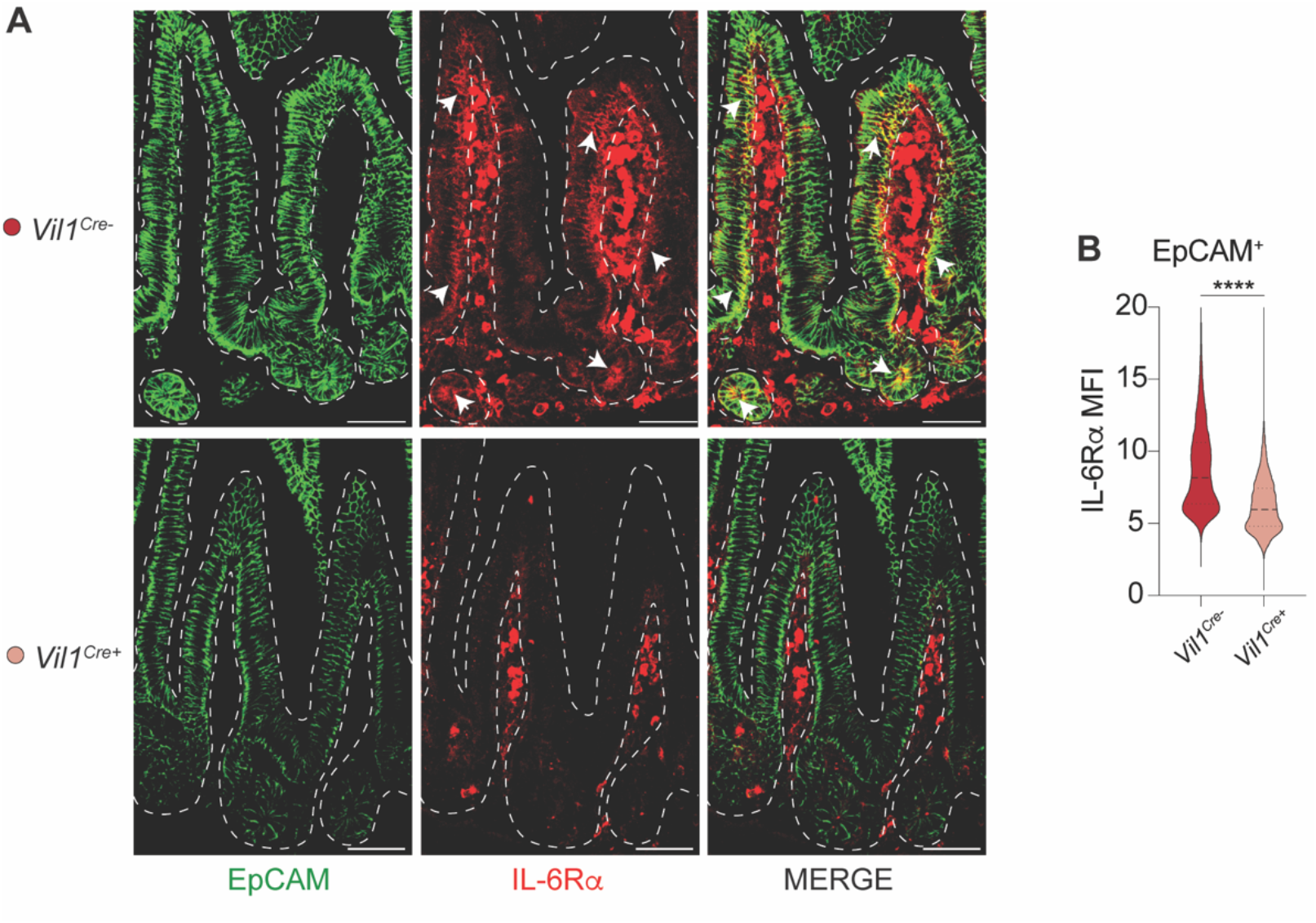
Specific *Il6ra* deletion from intestinal epithelial cells. (**A-B**) **(A**) Representative micrograph of small intestine from *Il6ra*^fl/fl^ *Vil1*^*Cre-*^ and *Il6ra*^fl/fl^ *Vil1*^*Cre+*^ offspring at 6 weeks-old stained for EpCAM and IL-6R*α*. Arrows point to IL-6R*α* expressed by EpCAM^+^ IEC (denoted by dashed line). Scale bars: 30 μm. **(B)** EpCAM^+^ IL-6R*α* MFI of small intestine of *Il6ra*^fl/fl^ *Vil1*^*Cre-*^ and *Il6ra*^fl/fl^ *Vil1*^*Cre+*^ offspring at 6 weeks-old. Data are representative of two independent experiments with 1-2 pregnant dams per group, using 2 offspring per pregnancy. **** p < 0.0001 (two-tailed unpaired Student’s t test).

**Figure S5.**
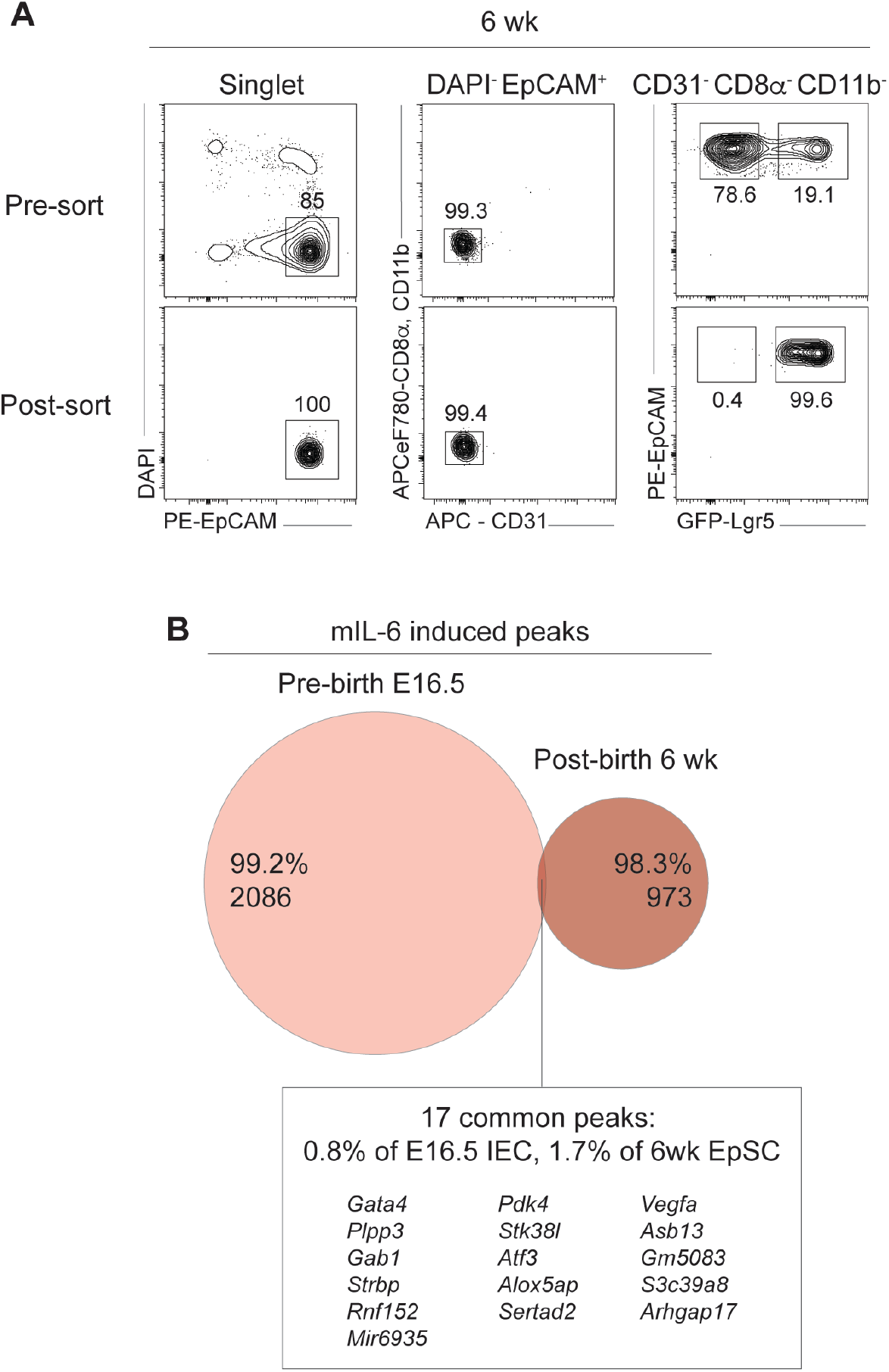
Increasing maternal IL-6 levels during pregnancy alters the chromatin accessibility of offspring intestinal epithelial stem cells. (**A**) Flow plots showing strategy to isolate intestinal epithelial stem cells (EpSC, EpCAM^+^ Lgr5^+^) and post-sort purity check. (**B**) Venn diagram shows the shared accessible ATAC-seq peaks unique to offspring from IL-6-injected dams (number indicated) between fetus IEC and adult EpSC. Data are from 1 experiment with 3 pregnant dams per group, using 3-7 offspring per pregnancy.

**Figure S6.**
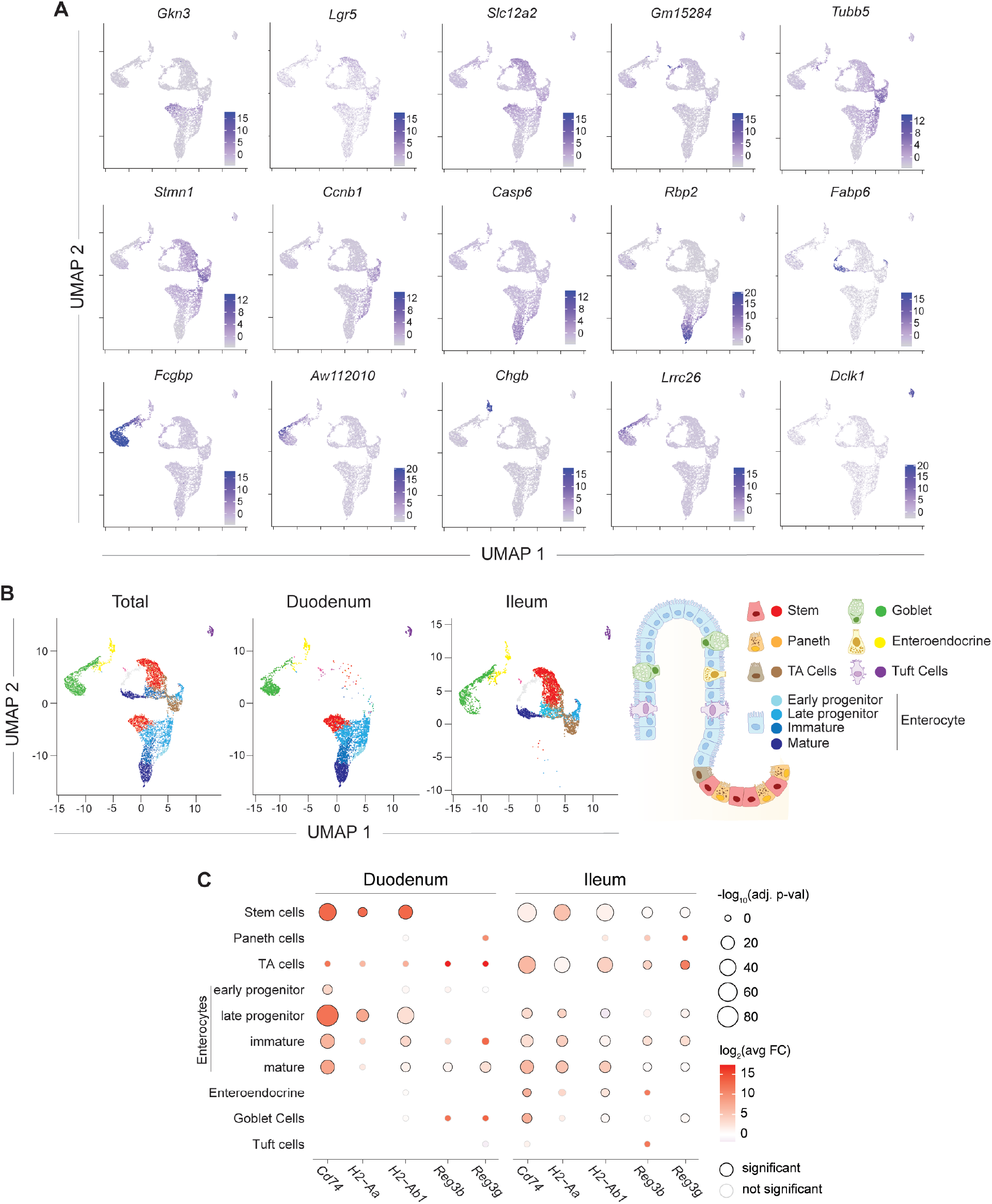
Increasing maternal IL-6 levels during pregnancy alters the transcriptome of offspring intestinal epithelial stem cells. **(A-C)** Intestinal epithelial cells (IEC) were sorted from mCtr or mIL-6 for scRNA-seq at 6 weeks old. (**A**) UMAP projection depicting transcriptional level of indicated genes enriched in specific cluster. (**B**) UMAP projection displaying distribution of IEC cluster from total samples, or IEC sorted from duodenum or ileum. Diagram illustrated different intestinal epithelial cell subsets. (**C**) Gene expression in epithelial cells subsets isolated from duodenum and ileum. Significance (dot size) and fold change (dot color) of differential gene expression by mIL-6 offspring compared to control offspring per IEC subsets. (**A-C**) Data are from 1 experiment with 3 pregnant dams per group, using 3-5 offspring per pregnancy.

**Figure S7.**
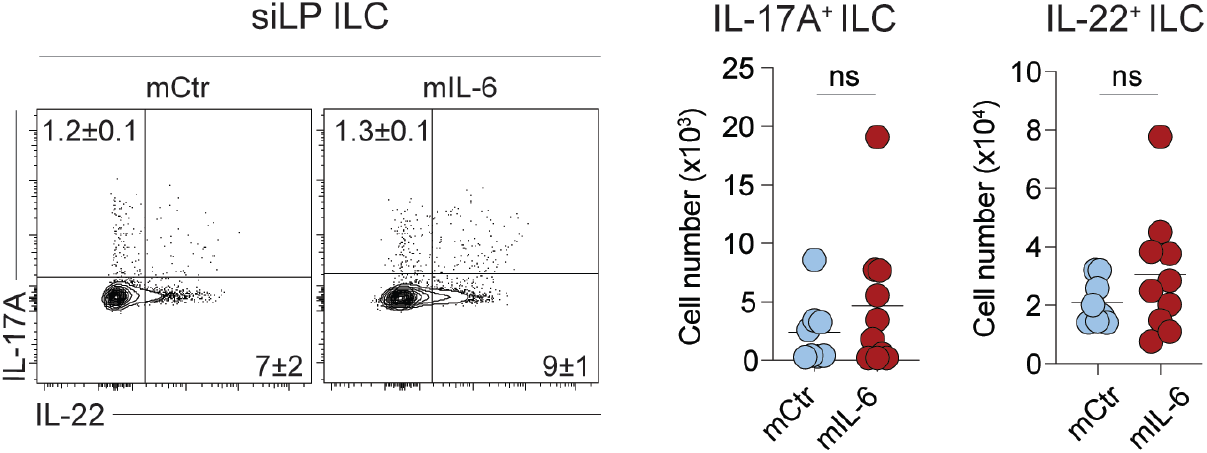
Increasing maternal IL-6 levels during pregnancy enhances protection to gastrointestinal infection in offspring. Timed-pregnant dams were injected with vehicle or 10 μg of IL-6 at gestation day 13.5 (E13.5), and their offspring (mCtr or mIL-6, respectively) were orally administered with *Salmonella* Typhimurim *(S*. Typhimurim) at 5 to 8 weeks old. Left: Representative contour plots showing cytokine production by siLP ILC (CD45^+^ CD90.2^+^ TCRβ^-^ TCRγδ^-^ CD127^+^). Right: Total IL-17A^+^ and IL-22^+^ siLP ILCs. Data are representative of 3 independent experiments with 1-2 pregnant dams per group, using 3-5 offspring per pregnancy. Numbers in representative flow plots indicate mean ± SD. Each dot represents an individual mouse. ns, not significant (two-tailed unpaired Student’s t test).

**Table S1.** Antibodies.

| **Antibodies** | **Sources** | **Identifier** |
| --- | --- | --- |
| Flow cytometry: anti-mouse CD45 (30-F11) APC-eFluor780 | Invitrogen | 47-0451-82 |
| Flow cytometry: anti-mouse CD90.2 (30-H12) BV785 | Biolegend | 105331 |
| Flow cytometry: anti-mouse TCR$\gamma\delta$ (GL3) BV650 | BD | 563993 |
| Flow cytometry: anti-mouse TCRb (H57-597) BUV737 | BD | 612821 |
| Flow cytometry: anti-mouse CD4 (RM4-5) BV510 | Biolegend | 100559 |
| Flow cytometry: anti-mouse CD8β (H35-17.2) BUV395 | BD | 740278 |
| Flow cytometry: anti-mouse CD8β (H35-17.2) PerCP-eFluor710 | Invitrogen | 46-0083-82 |
| Flow cytometry: anti-mouse Foxp3 (FJK-16s) AF700 | Invitrogen | 56-5773-62 |
| Flow cytometry: anti-mouse RORγt (Q331-378) | BD | 562607 |
| Flow cytometry: anti-mouse GATA3 (TWAJ) e660 | Invitrogen | 50-9966-42 |
| Flow cytometry: anti-mouse Tbet (4B10) BV421 | Biolegend | 644816 |
| Flow cytometry: anti-mouse IL17A (TC11-18H10.1) PeCy7 | Biolegend | 506922 |
| Flow cytometry: anti-mouse IFNγ (XMG1.2) BV605 | Biolegend | 505840 |
| Flow cytometry: anti-mouse IL-22 (1H8PWSR) PE | Invitrogen | 12-7221-82 |
| Flow cytometry: anti-mouse CD44 (IM7) AF700 | Invitrogen | 56-0441-82 |
| YopE:Kb+ Tetramer | NIH Tetramer Core | - |
| Phospho-Stat3 (Tyr703) (D3A7) XP Rabbit mAB (PE) | Cell Signaling | 8119S |
| Rabbit (DA1E) mAb IgG XP Isotype Control (PE) | Cell Signaling | 5742S |
| IEC sorting: EPCAM PE (G8.8) | Invitrogen | 12-5791-82 |
| IEC sorting: CD31 AF647 (MEC13.3) | Biolegend | 102516 |
| IEC sorting: CD8α APC-eFluor 780 (53-6.7) | Invitrogen | 47-0081-82 |
| IEC sorting: CD11b APC-eFluor (M1/70) | Invitrogen | 47-0112-82 |
| Immunohistochemistry: EPCAM-AF488 (G8.8) | Biolegend | 118210 |
| Immunohistochemistry: CD126-PE (D7715A7) | BD | 554462 |

## References

1. M. G. Netea et al., Defining trained immunity and its role in health and disease. Nat Rev Immunol 20, 375–388 (2020).

2. S. Naik et al., Inflammatory memory sensitizes skin epithelial stem cells to tissue damage. Nature 550, 475–480 (2017).

3. J. Ordovas-Montanes et al., Allergic inflammatory memory in human respiratory epithelial progenitor cells. Nature 560, 649–654 (2018).

4. R. S. Moore, R. Kaletsky, C. T. Murphy, Piwi/PRG-1 Argonaute and TGF-beta Mediate Transgenerational Learned Pathogenic Avoidance. Cell 177, 1827-1841 e1812 (2019).

5. G. Tetreau, J. Dhinaut, B. Gourbal, Y. Moret, Trans-generational Immune Priming in Invertebrates: Current Knowledge and Future Prospects. Front Immunol 10, 1938 (2019).

6. M. L. T. Berendsen et al., Maternal Priming: Bacillus Calmette-Guerin (BCG) Vaccine Scarring in Mothers Enhances the Survival of Their Child With a BCG Vaccine Scar. J Pediatric Infect Dis Soc 9, 166–172 (2020).

7. B. de Laval et al., C/EBPbeta-Dependent Epigenetic Memory Induces Trained Immunity in Hematopoietic Stem Cells. Cell Stem Cell 26, 657–674 e658 (2020).

8. S. P. Rosshart et al., Wild Mouse Gut Microbiota Promotes Host Fitness and Improves Disease Resistance. Cell 171, 1015–1028 e1013 (2017).

9. L. K. Beura et al., Normalizing the environment recapitulates adult human immune traits in laboratory mice. Nature 532, 512–516 (2016).

10. G. Mor, P. Aldo, A. B. Alvero, The unique immunological and microbial aspects of pregnancy. Nat Rev Immunol 17, 469–482 (2017).

11. S. Niewiesk, Maternal antibodies: clinical significance, mechanism of interference with immune responses, and possible vaccination strategies. Front Immunol 5, 446 (2014).

12. R. L. Goldenberg, J. C. Hauth, W. W. Andrews, Intrauterine infection and preterm delivery. N Engl J Med 342, 1500–1507 (2000).

13. R. Medzhitov, D. S. Schneider, M. P. Soares, Disease tolerance as a defense strategy. Science 335, 936–941 (2012).

14. J. L. McCarville, J. S. Ayres, Disease tolerance: concept and mechanisms. Curr Opin Immunol 50, 88–93 (2018).

15. S. J. Han et al., White Adipose Tissue Is a Reservoir for Memory T Cells and Promotes Protective Memory Responses to Infection. Immunity 47, 1154–1168 e1156 (2017).

16. K. Miyawaki et al., CD41 marks the initial myelo-erythroid lineage specification in adult mouse hematopoiesis: redefinition of murine common myeloid progenitor. Stem Cells 33, 976–987 (2015).

17. K. E. McGrath et al., Distinct Sources of Hematopoietic Progenitors Emerge before HSCs and Provide Functional Blood Cells in the Mammalian Embryo. Cell Rep 11, 1892– 1904 (2015).

18. Y. Cong, T. Feng, K. Fujihashi, T. R. Schoeb, C. O. Elson, A dominant, coordinated T regulatory cell-IgA response to the intestinal microbiota. Proc Natl Acad Sci U S A 106, 19256–19261 (2009).

19. T. W. Hand et al., Acute gastrointestinal infection induces long-lived microbiota-specific T cell responses. Science 337, 1553–1556 (2012).

20. T. Sano et al., An IL-23R/IL-22 Circuit Regulates Epithelial Serum Amyloid A to Promote Local Effector Th17 Responses. Cell 163, 381–393 (2015).

21. M. J. Molloy et al., Intraluminal containment of commensal outgrowth in the gut during infection-induced dysbiosis. Cell Host Microbe 14, 318–328 (2013).

22. V. J. Carrion et al., Pathogen-induced activation of disease-suppressive functions in the endophytic root microbiome. Science 366, 606–612 (2019).

23. M. V. Zaretsky, J. M. Alexander, W. Byrd, R. E. Bawdon, Transfer of inflammatory cytokines across the placenta. Obstet Gynecol 103, 546–550 (2004).

24. S. C. Ganal-Vonarburg, M. W. Hornef, A. J. Macpherson, Microbial-host molecular exchange and its functional consequences in early mammalian life. Science 368, 604– 607 (2020).

25. G. B. Choi et al., The maternal interleukin-17a pathway in mice promotes autism-like phenotypes in offspring. Science 351, 933–939 (2016).

26. S. Kim et al., Maternal gut bacteria promote neurodevelopmental abnormalities in mouse offspring. Nature 549, 528–532 (2017).

27. M. Murakami, D. Kamimura, T. Hirano, Pleiotropy and Specificity: Insights from the Interleukin 6 Family of Cytokines. Immunity 50, 812–831 (2019).

28. J. Dahlgren, A. M. Samuelsson, T. Jansson, A. Holmang, Interleukin-6 in the maternal circulation reaches the rat fetus in mid-gestation. Pediatr Res 60, 147–151 (2006).

29. L. Goetzl, T. Evans, J. Rivers, M. S. Suresh, E. Lieberman, Elevated maternal and fetal serum interleukin-6 levels are associated with epidural fever. Am J Obstet Gynecol 187, 834–838 (2002).

30. M. P. Verzi et al., Differentiation-specific histone modifications reveal dynamic chromatin interactions and partners for the intestinal transcription factor CDX2. Dev Cell 19, 713– 726 (2010).

31. U. Jadhav et al., Dynamic Reorganization of Chromatin Accessibility Signatures during Dedifferentiation of Secretory Precursors into Lgr5+ Intestinal Stem Cells. Cell Stem Cell 21, 65–77 e65 (2017).

32. R. Francis et al., Gastrointestinal transcription factors drive lineage-specific developmental programs in organ specification and cancer. Sci Adv 5, eaax8898 (2019).

33. N. Barker et al., Identification of stem cells in small intestine and colon by marker gene Lgr5. Nature 449, 1003–1007 (2007).

34. A. M. Chin, D. R. Hill, M. Aurora, J. R. Spence, Morphogenesis and maturation of the embryonic and postnatal intestine. Semin Cell Dev Biol 66, 81–93 (2017).

35. Z. Al Nabhani et al., A Weaning Reaction to Microbiota Is Required for Resistance to Immunopathologies in the Adult. Immunity 50, 1276–1288 e1275 (2019).

36. A. L. Haber et al., A single-cell survey of the small intestinal epithelium. Nature 551, 333– 339 (2017).

37. M. Biton et al., T Helper Cell Cytokines Modulate Intestinal Stem Cell Renewal and Differentiation. Cell 175, 1307–1320 e1322 (2018).

38. K. Atarashi et al., Th17 Cell Induction by Adhesion of Microbes to Intestinal Epithelial Cells. Cell 163, 367–380 (2015).

39. E. M. Velazquez et al., Endogenous Enterobacteriaceae underlie variation in susceptibility to Salmonella infection. Nat Microbiol 4, 1057–1064 (2019).

40. M. J. McGeachy, S. J. McSorley, Microbial-induced Th17: superhero or supervillain? J Immunol 189, 3285–3291 (2012).

41. K. Geddes et al., Identification of an innate T helper type 17 response to intestinal bacterial pathogens. Nat Med 17, 837–844 (2011).

42. S. Beyaz et al., High-fat diet enhances stemness and tumorigenicity of intestinal progenitors. Nature 531, 53–58 (2016).

43. L. M. Christian, K. Porter, Longitudinal changes in serum proinflammatory markers across pregnancy and postpartum: effects of maternal body mass index. Cytokine 70, 134–140 (2014).

## References

1. Y. Cong, T. Feng, K. Fujihashi, T. R. Schoeb, C. O. Elson, A dominant, coordinated T regulatory cell-IgA response to the intestinal microbiota. Proc Natl Acad Sci U S A 106, 19256–19261 (2009).

2. P. Shooshtari et al., Correlation analysis of intracellular and secreted cytokines via the generalized integrated mean fluorescence intensity. Cytometry A 77, 873–880 (2010).

3. E. Bolyen et al., Reproducible, interactive, scalable and extensible microbiome data science using QIIME 2. Nat Biotechnol 37, 852–857 (2019).

4. M. R. Corces et al., Lineage-specific and single-cell chromatin accessibility charts human hematopoiesis and leukemia evolution. Nat Genet 48, 1193–1203 (2016).

5. V. M. Link et al., Analysis of Genetically Diverse Macrophages Reveals Local and Domain-wide Mechanisms that Control Transcription Factor Binding and Function. Cell 173, 1796–1809 e1717 (2018).

6. M. Stoeckius et al., Cell Hashing with barcoded antibodies enables multiplexing and doublet detection for single cell genomics. Genome Biol 19, 224 (2018).

7. A. L. Haber et al., A single-cell survey of the small intestinal epithelium. Nature 551, 333– 339 (2017).

